# Coupling metabolic enhancement to plasmid spread enables programmable antimicrobial control

**DOI:** 10.64898/2026.02.25.707879

**Authors:** Pedro Dorado-Morales, Alicia Calvo-Villamañán, Bianca Audrain, Lisa Niedermeier, Morgan Lambérioux, Magaly Ducos-Galand, Bärbel Stecher, Jean-Marc Ghigo, Álvaro San Millán, Didier Mazel

## Abstract

The rise of multidrug-resistant pathogens underscores the need for precise antimicrobial strategies that extend beyond conventional antibiotics. Conjugation-based approaches offer a powerful yet underexploited means of delivering targeted genetic interventions directly within microbial communities. In this work, we combined selective killing modules with rationally optimized conjugative vectors to target antibiotic-resistant pathogens and clinically relevant antimicrobial resistance plasmids. First, we engineered and validated toxin-intein modules, programmable cassettes that restrict toxic activity to highly specific regulatory contexts. Specifically, we developed and validated modules targeting *Shigella* spp., *Salmonella enterica*, and bacteria carrying the resistance plasmid pOXA-48, demonstrating a tunable system capable of selective activity at both the species and strain levels. To identify the most effective delivery platform, we compared mobilizable and conjugative systems and found that, *in vitro*, conjugative plasmids consistently outperformed mobilizable ones by approximately one order of magnitude. To further optimize delivery, we streamlined the broad-host-range plasmid RP4 and enhanced its functionality by incorporating either the metabolic *fos* locus, which confers a fitness advantage to cells carrying the delivery vehicle; a type IV pilus operon that promotes mating-pair stabilization and enables efficient conjugation in liquid environments; or both features combined. Using these engineered RP4 derivatives, we integrated the toxin–intein module targeting pOXA-48 and evaluated its performance in complex microbial communities. In this setting, the RP4 variant carrying both the *fos* locus and the type IV pilus operon effectively blocked the spread of pOXA-48. Together, this work advances the use of conjugative plasmids as robust and programmable platforms to combat antibiotic resistance and enable microbiome engineering. Beyond introducing highly specific antimicrobial modules and a new generation of optimized conjugative vectors, our results identify ecological competitiveness and plasmid transfer dynamics as critical determinants of the success of such interventions.

## Introduction

In recent decades, the widespread and often indiscriminate use of antimicrobial agents, particularly in clinical and agricultural settings, has led to the emergence and proliferation of multidrug-resistant pathogens^1^. The growing <u>a</u>nti<u>m</u>icrobial resistance (AMR) crisis is further complicated by the broad-spectrum activity of most conventional antibiotics, which not only act on target bacteria but also affect other components of the microbial community^2^. This disruption not only results in ecological imbalances, but also facilitates horizontal transfer of resistance genes, accelerating the evolution and acquisition of AMR mechanisms^3^. As antibiotics are one of the cornerstones of modern medicine, their decreasing efficacy poses one of the greatest threats to public health. In fact, annually, in the European Union alone, AMR causes around 35,000 deaths^4^ and imposes an estimated economic burden of 11.7 billion euros^5^.

To address this issue, in 2024, the <u>W</u>orld <u>H</u>ealth <u>O</u>rganization (WHO) updated its list of priority pathogens to orient the focus of research and development of new effective antimicrobial treatments against resistant bacteria^6,7^. Among the urgent threats are fluoroquinolone-resistant *Salmonella* and *Shigella* species^6,8,9^. *Salmonella enterica* is estimated to cause 14 million infections and 135,000 deaths globally each year^10^, while *Shigella* was reported as the second leading cause of diarrhoeal mortality in 2016, being responsible for over 212,000 deaths^11^. In addition to these pathogens, carbapenem-resistant *Enterobacteriaceae* are classified by the WHO as critical-priority pathogens due to their limited treatment options and high mortality rates^7^.

Conjugative plasmids are the main vehicles for the dissemination of carbapenem and other AMR genes in clinical settings and, as such, represent critical targets for intervention. Many of these plasmids have a broad host range that allows for the rapid dissemination of AMR genes across bacterial species in the gut^12^. Amongst them, pOXA-48 stands out as a highly prevalent carbapenem-resistance plasmid whose spread must be contained. pOXA-48 was first identified in a *Klebsiella pneumoniae* clinical isolate in Turkey in 2001^13^ and is currently associated with carbapenemase-producing *Enterobacteriaceae* outbreaks worldwide^14^. Carbapenem antibiotics are used in hospitals as last-resort drugs to treat multidrug-resistant infections. Therefore, developing strategies to prevent the dissemination of plasmids such as pOXA-48 is of critical importance.

Despite the need for solutions, the discovery and development of new classes of antibiotics has been in constant decline for years. As a result, alternative approaches, such as the use of monoclonal antibodies^15^, antimicrobial peptides^16^, bacteriophages^17,18^, lysins^19,20^ and CRISPR-Cas-based technologies^21^, are receiving increasing attention. While many of these strategies show promise in terms of precision and specificity, challenges remain in terms of delivery, host range and the risk of resistance development.

Previous work by two of the groups involved in this study demonstrated that genetic modules based on split toxins reconstituted through intein-mediated protein splicing, and delivered via conjugation, can serve as highly effective antimicrobial agents^22^. In these systems, each half of a toxin is fused to one half of a split intein, and expression of the toxin–intein constructs is driven by transcription factors specific to the target organism. A key advantage of this strategy is the additional layer of post-translational regulation provided by the intein, which prevents the reconstitution of an active toxin in non-target species.

Inteins are self-catalytic protein segments capable of excising from a larger precursor protein in which they are embedded. This leads to the ligation of the protein sequences flanking the intein through the formation of a new peptide bond^23^. Inteins can differ in their splicing efficiency, a property that can be exploited to suppress basal expression from leaky promoters and ensure that toxin reconstitution occurs only when expression levels exceed a defined threshold^24^.

This strategy of conditional toxin expression and reconstitution via intein-mediated splicing proved to be highly modular and adaptable, as illustrated by its successful application in selectively eliminating either all *Vibrio cholerae* cells or specifically their antibiotic-resistant derivatives from mixed bacterial populations^22^.

Building on these findings, this study seeks to broaden the applicability and toolkit of this technology as a step toward the development of the next generation of targeted antimicrobials. Specifically, we sought to: (1) develop toxin–intein modules to target priority pathogens and clinically relevant AMR mobile genetic elements; (2) identify and optimize the delivery platform for the dissemination of antibacterial modules; (3) develop engineered derivatives of the delivery system with enhanced functionality in complex microbial populations; and (4) validate system performance *in vitro*, both in defined conjugation assays and in complex microbial communities.

## Results

### 1. Rewiring toxin–intein modules to target priority pathogens and carriers of AMR plasmids

To broaden the scope of toxin–intein antimicrobials beyond *Vibrio cholerae*^22^, we engineered modules targeting two WHO priority pathogens, *Shigella* and *Salmonella enterica*, as well as cells carrying the clinically relevant carbapenem-resistance plasmid pOXA-48.

All systems were designed using a minimal architecture in which the expression of a toxin-intein pair is driven by a single, target-specific transcriptional activator (**Figure S1**). For *Shigella* and *Salmonella*, we selected regulators associated with genus-specific virulence loci^25,26^, whereas for pOXA-48 we exploited a plasmid-encoded regulator involved in plasmid–chromosome transcriptional crosstalk^27^.

#### 1.1. Targeting the VirF regulatory axis selectively eliminates *Shigella*

From a taxonomic perspective, all members of the *Shigella* genus are considered part of the broader *Escherichia* genus. What primarily distinguishes *Shigella* species and <u>e</u>nteroinvasive *<u>E</u>. <u>c</u>oli* (EI*EC*) from other *Escherichia* strains is the presence of the virulence plasmid pINV. This large (210–230 kb), non-self-transmissible plasmid is essential for the invasive phenotype that characterizes both *Shigella* and EI*EC*^28^.

Virulence in these pathogens is mainly regulated by VirF, a DNA-binding transcriptional activator belonging to the AraC family. VirF plays a central role in initiating a regulatory cascade that enables bacterial invasion. It activates the transcription of key virulence genes, including *virB* and *icsA*, by binding directly to their promoter regions. *virB* encodes a secondary regulator that further drives the expression of genes required for the assembly of a type III secretion system (T3SS) that facilitates invasion of epithelial cells^25,29^ (**Figure S2A**).

Given its central regulatory role, VirF represents a promising anti-virulence target. For this reason, we sought to exploit this regulatory circuit as the foundation of our anti-*Shigella*/EI*EC* strategy. As a first step, we evaluated the activity of the *virB* promoter, aiming to assess its responsiveness to VirF and determine its suitability for driving expression of a toxin in a VirF-dependent system (**Supplementary information 1**).

Right after, the toxin–intein module described by Rocío López et *al*.^22^, consisting on the gene encoding the gyrase inhibitor CcdB^30^ interrupted by the fast-splicing iDnaE^31,32^ intein–coding sequence (*ccdB*::*iDnaE*), was cloned under the control of the *virB* promoter (**Figure S2C**) into a mobilizable, low-copy-number plasmid. The resulting construction was then used to evaluate the anti-*Shigella*/EI*EC* system.

Following conjugation and selection of transconjugants, no toxic effect was observed at 30°C in any of the recipient strains, including those harboring pINV and the negative control *E. coli* MG1655. In contrast, at 37°C, a marked reduction in the number of recovered transconjugants was observed across all *Shigella* species tested (**Figure 1A**). This outcome is consistent with the temperature-dependent activation of the *virB* promoter via VirF (**Figure S3A**).

**Figure 1.**
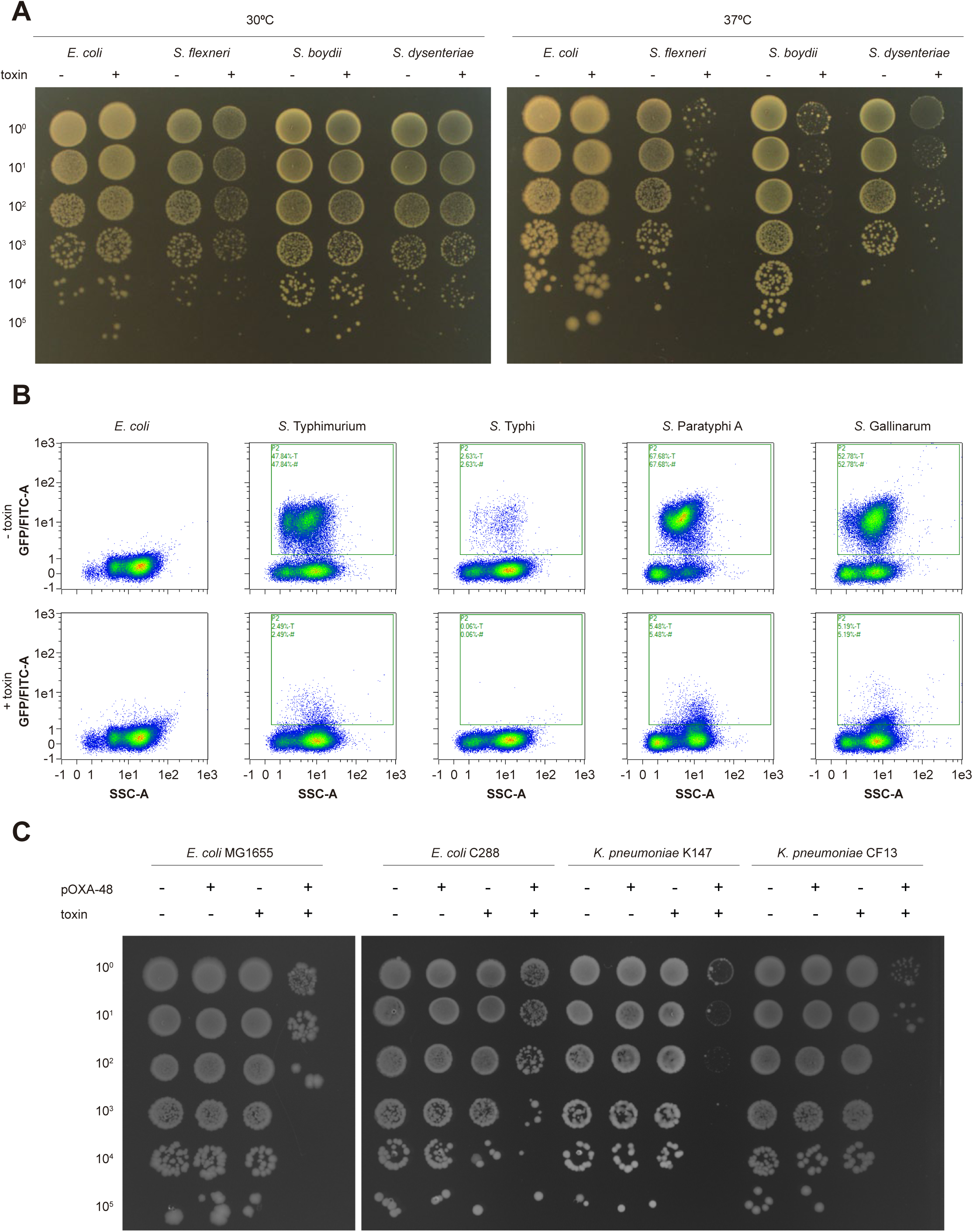
Validation of toxin-intein modules. **A.** Delivery of the *virF*-based toxin-intein module into *E. coli* MG1655 (control) and three *Shigella* species (*S. flexneri*, *S. boydii* and *S. dysenteriae*). Recipient strains were transferred either the empty plasmid or its derivative carrying the toxin-intein system. Transconjugants obtained from experiments at 30°C and 37°C are shown in the left and right panel, respectively. **B.** Validation of the *Salmonella enterica* toxin-intein module. *E. coli* (control) and various *S. enterica* strains (Typhimurium, Typhi, Paratyphi A and Gallinarum), all carrying a compatible *hilA* reporter plasmid, were transferred the empty vector (upper panel) or the same plasmid carrying the toxin-intein cassette (lower panel). Reporter fluorescence was monitored in transconjugants by flow cytometry under *hilD*-inducing conditions. Data correspond to the 8-hour time point for all strains except for Gallinarum, for which the 24-hour time point is shown due to its slower growth. **C.** Delivery of the anti-pOXA-48 toxin–intein module into *E. coli* MG1655 and three clinical enterobacterial isolates (*E. coli* C288, *K. pneumoniae* K147 and *C. freundii* CF13). Recipient strains, either carrying pOXA-48 or plasmid-free, received either the empty delivery plasmid or its derivative encoding the anti-pOXA-48 toxin–intein system. The data shown in this figure are representative of at least three independent biological replicates.

Despite this selective effect, a subset of surviving transconjugants, hereafter referred to as “escapers”, was still detected at varying frequences depending on the species. To investigate the underlying mechanisms, we focused on *Shigella flexneri* and screened 240 escaper clones by PCR for three regions: *virF*, *virB* and the toxin-intein module. Among these clones, one harbored a mutation within the toxin-intein cassette caused by the insertion of an IS1F element in the N-terminal part of the intein. Eleven additional clones retained *virB* but lacked *virF*, likely due to IS-mediated pINV rearrangements. The remaining clones tested negative for both *virF* and *virB* while still retaining the toxin-intein module. These results suggest either loss of the pINV plasmid or large-scale rearrangements leading to deletion of extensive plasmid regions.

In conclusion, the toxin-intein system performs as intended and demonstrates specificity toward *Shigella* and, presumably, EI*EC*. The vast majority of escapers result from the loss of the *virB* and *virF* sequences which likely renders them avirulent. However, a small proportion of escape events appears to be associated with mutations within the toxin-intein module itself. Further investigation will be needed to determine whether the frequency of such escapers is sufficient to allow *in vivo* colonization and the development of shigellosis.

#### 1.2. A HilD-responsive toxin–intein module blocks *Salmonella* virulence activation

*Salmonella enterica* relies on Salmonella Pathogenicity Island 1 (SPI-1) for the invasion of intestinal epithelial cells, a key step in establishing infection^33^. The expression of SPI-1 genes is orchestrated by a complex regulatory network, with HilD, an AraC-like transcriptional regulator, as master regulator^34^. HilD responds to environmental cues associated with the intestinal microenvironment, directly activating the transcription of genes within and outside SPI-1^35^. Among its direct targets is *hilA*, which encodes a transcription factor of the OmpR/ToxR family responsible for activating the genes involved in the assembly of a functional T3SS that is essential for bacterial invasion^36–40^ (**Figure S4A**). Additionally, as demonstrated by Petrone *et al*.^26^, the *hilA* promoter shows minimal basal activity in the absence of HilD, making it an attractive candidate for driving the expression of our synthetic systems.

After confirming that the *hilA* promoter met our criteria for expression strength and specificity using a reporter system analogous to the one previously developed for *Shigella* (**Supplementary information 2**), we moved on to design the corresponding anti-*Salmonella* strategy. As targeting virulence rather than bacterial viability is thought to exert weaker selective pressure for the evolution of resistance mechanisms^41^, we decided to explore a strategy distinct from the one used for *Shigella*. Specifically, we employed a toxin-intein module composed of the toxin HigB2^42^ and the intein iGyrB^43^ (**Figure S4D**). In this system, HigB2, a ribosome-dependent RNase that cleaves translating mRNAs^44^, induces delayed toxicity rather than immediate cell death, as shown in Figure S6 of López-Igual *et al*.^45^ This delay in the cytotoxic effect raises the possibility that, upon induction of *hilD*, HigB2 could degrade actively translated mRNAs, effectively “resetting” the cell’s virulence program. Moreover, by optimizing the translation initiation of HigB2 through replacement of *hilA* native Shine-Dalgarno sequence with a stronger consensus motif (**Supplementary information 2**), toxin production is expected to exceed that of HilA, enhancing the robustness of the system.

Since we could not monitor a loss-of-viability phenotype, we instead followed the dynamics of *hilA* expression by introducing the reporter described in **Supplementary Information 2** into multiple *Salmonella enterica* strains. In parallel, the *higB2*::*iGyrB* toxin-intein cassette was placed under the control of the *hilA* promoter, together with the modified 5′ UTR designed to enhance translation initiation, and cloned into a mobilizable, low-copy-number plasmid. Conjugation assays were then performed, and reporter activity was monitored under *hilD*-inducing conditions^46–48^ in transconjugants carrying either the empty vector or the toxin–intein system.

As shown in **Figure 1B** and **S5**, all tested *Salmonella* strains exhibited a similar pattern: in the presence of the empty plasmid, a subpopulation of cells, varying in size depending on the strain, activated *hilA* expression. In contrast, when the toxin-intein module was delivered, the emergence of this *hilA*-expressing subpopulation was markedly impaired, indicating effective inhibition of *hilA* activation by the toxin system.

Altogether, these results demonstrate that conditional deployment of a toxin–intein module targeting the HilD regulatory axis effectively disrupts virulence program activation across multiple *Salmonella enterica* serovars. Rather than inducing rapid cell death, this strategy interferes with the transcriptional cascade required for T3SS expression. Importantly, this mode of action suggests that virulence blockade can be as effective as bactericidal approaches while potentially reducing selective pressure for resistance and the emergence of escapers, consistent with previous reports highlighting the robustness of inhibitory strategies such as <u>CRISPR i</u>nterference (CRISPRi)^49^.

#### 1.3. Exploiting a plasmid–chromosome crosstalk allows selective targeting of pOXA-48 carriers

##### 1.3.1. Plasmid–chromosome transcriptional crosstalk mediated by antagonistic LysR regulators enables sensing of pOXA-48

Finally, we sought to determine whether our toxin-intein-based strategy could be extended to selectively eliminate bacteria carrying clinically relevant AMR plasmids. As a proof of concept, we focused on pOXA-48, a family of conjugative plasmids encoding the carbapenemase gene *blaOXA-*48.

To this end, we exploited a recently described plasmid–chromosome transcriptional crosstalk mediated by a LysR-type transcriptional regulator encoded on pOXA-48 (hereafter referred to as LysR_pOXA-48_)^27^. This regulator activates the chromosomal *pfp*–*ifp* operon, whose promoter can therefore act as a sensor for the presence of pOXA-48 (**Figure S6A**).

The *pfp*–*ifp* operon is widely conserved among *Enterobacteriaceae* associated with pOXA-48 carriage (e.g. >84% of *Klebsiella* spp., >78% of *Citrobacter* spp. and >99% of *Salmonella* spp.), with the notable exception of *Escherichia coli*, in which this operon is virtually absent.

The *pfp*–*ifp* operon is preceded by a LysR-type transcriptional regulator encoded upstream and divergently oriented relative to the operon (hereafter referred to as LysR_chr_). To assess potential regulatory interactions between the *pfp*–*ifp* promoter, LysR_chr_ and the plasmid-encoded LysR (LysR_pOXA-48_), we constructed two transcriptional reporter systems. In the first reporter, *gfp* expression was placed under the control of the *pfp*–*ifp* promoter together with an intact copy of *lysR_chr_* (**Figure S6B**). In the second construct, the same promoter–*gfp* fusion was preceded by a methionine-less version of *lysR_chr_*, thereby preventing its translation (**Figure S6C**).

Consistent with previous reports^27^, we observed that the *pfp*–*ifp* promoter was activated in the presence of pOXA-48, confirming that LysR_pOXA-48_ functions as an activator of this promoter (**Figure S3C**). In contrast, impairing *lysR_chr_* translation resulted in increased promoter activity, consistent with a potential repressive role of LysR_chr_ on the *pfp*–*ifp* operon (**Figure S6A**). However, additional experiments will be required to fully disentangle this regulatory interaction, as we cannot exclude the possibility that removal of the *lysR_chr_* start codon also affects nearby regulatory DNA elements involved in LysR_chr/pOXA-48_ binding.

##### 1.3.2. Conjugative delivery of a toxin–intein module enables selective elimination of pOXA-48 carriers

Based on the results described above, we selected the configuration in which *lysR_chr_* is not translated as the most responsive and specific sensor for pOXA-48 (**Figure S3C**). For the toxin–intein module, we employed the gyrase inhibitor ParE2^50^ together with the fast-splicing intein iDnaE. Accordingly, the *parE2*::*iDnaE* module^45^ was placed under the control of the *pfp*–*ifp* promoter in the methionine-less *lysR_chr_* configuration (**Figure S6D**) and cloned into a mobilizable, low-copy-number plasmid. The iDnaE intein is characterised by high splicing efficiency^31,32^, which initially resulted in toxicity and poor recovery of transformants. Among those, we were able to isolate intein variants carrying mutations that partially reduced intein splicing efficiency, yielding toxin–intein modules that were compatible with host viability but remained functional upon activation.

The resulting vectors were introduced into the donor strain NGE*pir* to enable mobilisation, and conjugation assays were performed as described in the Methods section. Delivery of the anti-pOXA-48 toxin–intein module resulted in efficient and highly specific killing of clinical *Enterobacteriaceae* strains carrying pOXA-48, while plasmid-free control strains remained unaffected (**Figure 1C**).

As observed with the anti-*Shigella* modules, a small fraction of surviving transconjugants carrying both pOXA-48 and the anti-pOXA-48 vector emerged, with frequencies that varied across species. We analysed twelve escaper clones isolated from all tested species. Among these, six carried inactivating mutations in the *parE2*::*iDnaE* killing cassette, three harboured mutations in *lysR_pOXA-48_*, and two contained mutations in both elements. Only a single escaper retained intact versions of both *lysR_pOXA-48_* and the *parE2*::*iDnaE* module.

Together, these results demonstrate that the anti-pOXA-48 system enables highly specific discrimination between pOXA-48–carrying and plasmid-free bacteria, leading to efficient elimination of resistance plasmid carriers across multiple clinically relevant *Enterobacteriaceae* species.

### 2. RP4 achieves higher transfer efficiency than mobilisable vectors

Once we had generated the new toxin–intein modules, we next sought to determine the most effective plasmid configuration for their delivery. Specifically, we compared the performance and dissemination potential of the three most commonly used setups in the literature: a mobilizable plasmid lacking self-transmissibility and relying on a conjugative helper^22,51^; a mobilizable plasmid parasitising a fully functional conjugative element^52^; and a single, fully self-transmissible conjugative plasmid^53,54^ (**Figure S7**).

To enable a direct experimental comparison of these strategies, we reconstructed each configuration in an otherwise identical genetic background. In the first setup, the mobilizable plasmid pRC2^55^ was co-introduced with a streamlined version of RP4 carrying a 15-nucleotide deletion in the *oriT* sequence^56^ (stRP4 Δ35), a modification that abolishes RP4 transfer while preserving its ability to function as a conjugative helper (**Figure S7A**). In the second setup, the same mobilizable plasmid pRC2 was paired with the streamlined RP4 variant (stRP4) containing an intact *oriT*, allowing both plasmids to be mobilised during conjugation (**Figure S7B**). Finally, in the third configuration, the stRP4 plasmid was used alone, functioning as a fully self-transmissible element (**Figure S7C**).

We selected pRC2 as the mobilizable vector rather than a plasmid carrying the RP4 *oriT* for two key reasons. First, pRC2 encodes its own *oriT* and cognate relaxase, which are fully compatible with the RP4-encoded type <u>IV</u> <u>s</u>ecretion <u>s</u>ystem (T4SS). Second, pRC2 shares no sequence homology with RP4, thereby minimizing the risk of homologous recombination and unintended co-transfer events (plasmid conduction). In parallel, we employed a streamlined version of RP4 (stRP4; **Supplementary Information 3**) to reduce the likelihood of of unintended integration or rearrangement events, such as those mediated by insertion sequences.

Having established these three configurations, we next compared their transfer efficiencies *in vitro*. Conjugation assays were performed using an auxotrophic *E. coli* strain as the donor, allowing efficient counterselection, and its wild-type counterpart as the recipient. As shown in **Figure 2**, the stRP4 plasmid, either alone (setup 3) or coexisting with the mobilizable plasmid (setup 2), was able to fully invade the recipient population, with transfer frequencies consistently above 80% and reaching up to ∼99% across replicates (mean 0.93 ± 0.06; n = 8). In contrast, the mobilizable plasmid pRC2 displayed an approximately tenfold lower transfer efficiency, regardless of whether it was paired with a non-transmissible or a fully conjugative RP4 helper (setups 1 and 2, respectively). Notably, no significant difference was observed between these two conditions, indicating that the presence of a functional conjugative plasmid does not measurably enhance mobilizable plasmid dissemination under the conditions tested.

**Figure 2.**
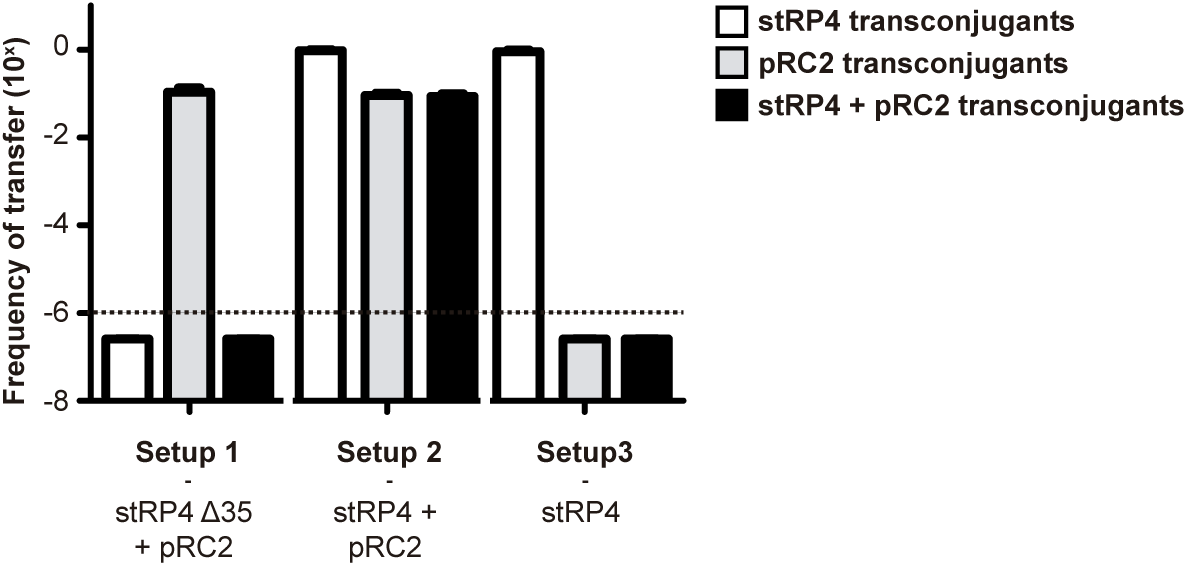
Transfer efficiencies across different plasmid configurations. **A.** Transfer frequencies of the streamlined RP4 (stRP4), the mobilisable plasmid pRC2, and both plasmids together were measured in solid-surface mating assays with *E. coli* MG1655 as recipient. White bars indicate the frequency of stRP4 transfer; grey bars correspond to pRC2 transfer; and black bars represent co-transfer of both plasmids. Data are shown as mean values with standard deviations from four independent biological replicates. The dashed line marks the detection limit.

Based on these results, we focused exclusively on conjugative plasmids for subsequent experiments. Although conjugative plasmids are intrinsically more complex than mobilisable vectors, RP4 represents a well-characterized model system, whose genetic organization and regulatory elements have been extensively studied^57–61^, facilitating rational engineering despite its size.

### 3. Rational engineering of RP4 improves both plasmid fitness and conjugation efficiency

*In vitro* assays demonstrated that the conjugative plasmid consistently outperformed both mobilizable vector configurations. Nevertheless, because the gut represents a heterogeneous and highly competitive environment, we therefore sought to enhance stRP4 performance by incorporating features aimed at improving plasmid transfer efficiency and host competitiveness. To this end, we constructed three derivative versions using the previously described stRP4 as a backbone, each specifically designed to address these constraints.

#### 3.1. The *fos* locus confers a competitive fitness advantage to stRP4-bearing bacteria

The first version was designed to provide a positive selective advantage to RP4-bearing bacteria. Given that <u>m</u>obile <u>g</u>enetic <u>e</u>lements (MGEs) often carry adaptative traits, we searched for *loci* found in such elements that had also been shown to provide functional advantages and that were not linked to virulence nor antimicrobial resistance. One such candidate was the *fos* locus, which has been associated with enhanced bacterial growth and improved colonization in the gut environment^62^. This locus comprises six genes organized as an operon (*fosK*, *fosY*, *fosGH2*, *fosX*, *fosGH1* and *fosT*), along with a divergently transcribed gene (*fosR*) encoding a putative repressor of the operon^62,63^. The operon encodes a sugar transporter and several enzymes responsible for the metabolism of <u>s</u>hort <u>c</u>hain fructo<u>o</u>ligo<u>s</u>accharides (scFOS), fructose oligomers composed of a terminal glucose residue and two to four fructose units. These compounds are not hydrolysed by host digestive enzymes and therefore reach the intestine intact, where they can be used as a carbon source by probiotic bacteria and strains harbouring the *fos* locus.

After having cloned the *fos* locus into the reduced version of the RP4 plasmid and confirmed its functionality (**Supplementary information 4**), we assessed whether carrying the *fos* operon conferred a competitive advantage under different nutrient conditions. To do this, we performed competition assays in the presence and absence of exogenous scFOS. To distinguish between the two plasmids, we generated alternative versions of both, the stRP4 and its *fos* derivative, in which the *aphA* gene (kanamycin resistance) was replaced with the *bla* gene (resistance to β-lactam antibiotics). These *bla*-carrying plasmids will be referred from here on as “AmpR”. As shown in **Figure 3A**, when comparing strains carrying stRP4 and stRP4 AmpR, the strain harbouring the AmpR version exhibited a fitness defect, likely due to periplasmic stress caused by the accumulation of β-lactamase. Interestingly, this burden was mitigated by the presence of the *fos* locus, even in the absence of exogenously added scFOS (**Figure 3B** and **D**, **left panels**). This finding is unexpected, as the *fos* operon is theoretically repressed under these conditions^63^.

**Figure 3.**
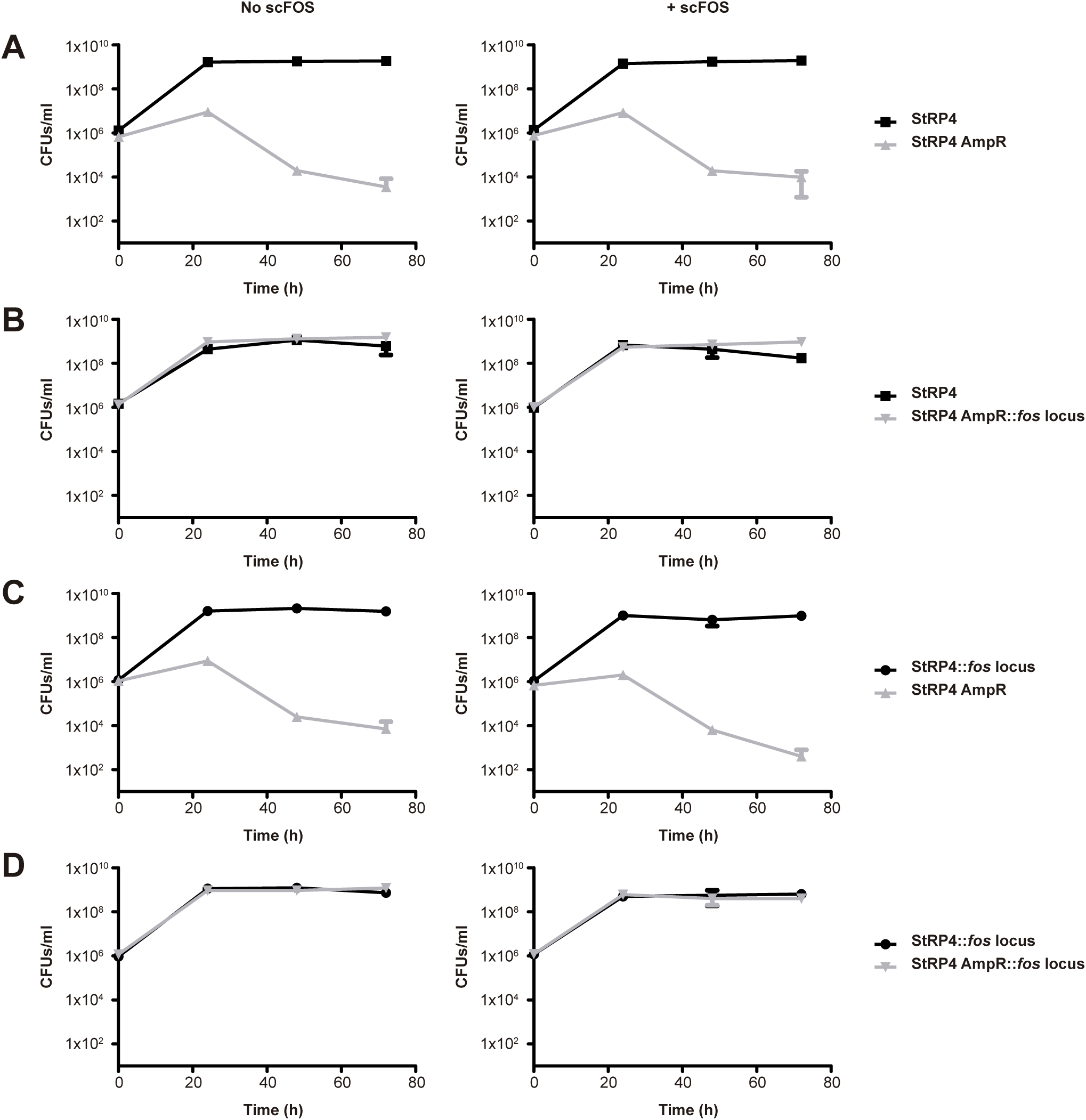
Effect of the *fos* locus on *E. coli* MG1655 fitness. Competition assays between isogenic *E. coli* MG1655 strains carrying different stRP4 variants, performed in the absence (left) and presence of scFOS (0.5% w/v) (right). Top panel: stRP4 (black squares) vs. stRP4 AmpR (grey triangles). Second panel: stRP4 (black squares) vs. stRP4 AmpR::*fos* locus (inverted grey triangles). Third panel: stRP4::*fos* locus (black circles) vs. stRP4 AmpR (grey triangles). Bottom panel: stRP4::*fos* locus (black circles) vs. stRP4 AmpR::*fos* locus (inverted grey triangles). Data represent mean values from three independent biological replicates. Error bars indicate standard deviation.

Nevertheless, as expected, the competitive advantage conferred by the *fos* locus is markedly enhanced in the presence of scFOS, as evidenced by the approximate tenfold reduction in the population of cells carrying the stRP4 AmpR plasmid when competed with their isogenic counterparts containing the stRP4 plasmid that carries the *fos* locus (**Figure 3C**).

Altogether, these results indicate that the *fos* locus confers a measurable fitness advantage to stRP4-bearing bacteria, improving plasmid persistence and partially offsetting plasmid-associated costs. These properties make the *fos* locus an attractive trait for stabilizing conjugative delivery platforms in competitive and nutrient-variable environments such as the gut.

#### 3.2. Equipping stRP4 with a type IV pilus overcomes its limitation in liquid conjugation

Conjugative pili involved in plasmid transfer can be broadly classified into two types based on their structure and function: long, flexible pili and short, rigid pili^64^. Plasmids encoding long, flexible pili, such as those from the IncF, IncH and IncI groups, are capable of efficient conjugative transfer in both liquid and solid environments. In contrast, plasmids that produce short, rigid pili, such as those of the IncN, IncP and IncW incompatibility groups, conjugate with higher efficiency (typically 2 to 4 orders of magnitude) on solid surfaces compared to liquid media. This disparity is largely attributed to the ability of flexible pili to mediate mating pair stabilization^65^, a feature that also appears to be important for efficient conjugation within the gut microbiota^66^.

RP4 encodes rigid pili and therefore displays reduced conjugation efficiency in liquid compared to solid environments. Certain plasmids, such as R64, have been described to encode both types of pili. In the case of R64, the thin pilus, classified as a type IV pilus, is essential for conjugative transfer in liquid conditions. Fourteen genes (*pilI* through *pilV*) are involved in the formation of the thin pilus, with the *pilV* gene being particularly noteworthy due to the presence of a variable C-terminal domain encoded by a DNA rearrangement system known as shufflon^67^. This system consists of multiple invertible DNA segments flanked by recombination sites that can be rearranged by a shufflon-specific tyrosine recombinase encoded by the neighbouring *rci* gene. The various PilV C-terminal variants produced through shufflon-mediated recombination act as adhesins that modify plasmid’s host range by altering recipient cell recognition^68^. Additionally, the *traABCD* regulatory cluster, located upstream of the type IV pilus operon, plays a key role in regulating thin pilus formation and conjugative transfer in both liquid and solid environments, with *traB* and *traC* acting as positive regulators of transfer gene expression^69^.

With this in mind, we cloned the entire *pil* operon, including the shufflon region and the *rci* recombinase gene, and the *traABCD* regulatory cluster from the derepressed conjugative plasmid R64drd11 into the stRP4 backbone. The goal was to enhance the conjugation efficiency of stRP4 in liquid environments by equipping it with a complete and functional type IV pilus.

To assess whether the incorporation of the R64drd11 type IV pilus improved the conjugation efficiency of stRP4, we evaluated plasmid transfer under both solid and liquid mating conditions across different recipient species: *E. coli*, *S. flexneri* and *S. enterica*.

As shown in **Figure 4A**, under solid mating conditions, the addition of the type IV pilus had no measurable impact on the conjugation efficiency when the plasmid was transferred from *E. coli* to either *E. coli* or *Shigella flexneri*. However, its transfer from *E. coli* to *Salmonella enterica* was negatively affected compared to stRP4, displaying a reduction of approximately one order of magnitude.

**Figure 4.**
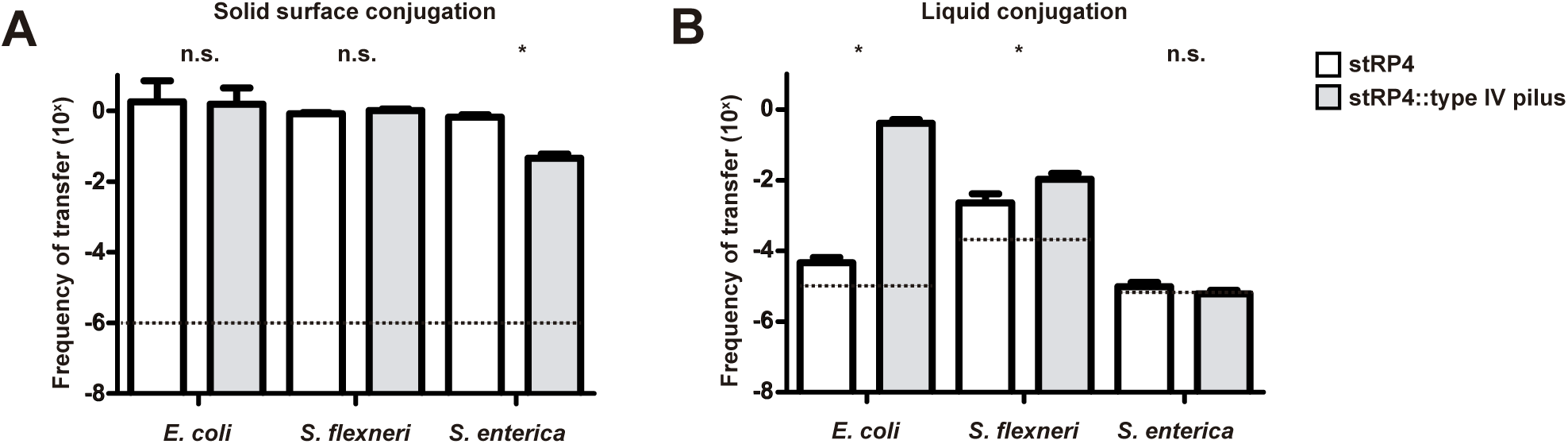
Comparison of stRP4 (white) and stRP4::type IV pilus (grey) transfer efficiencies using solid surface (A) and liquid conjugation (B) assays. An auxotrophic *E. coli* strain was used as donor and *E. coli* MG1655, *Shigella flexneri* 2a or *Salmonella enterica* SL1344 as recipients. Data represent mean values from four independent biological replicates. Error bars indicate standard deviation. Statistical analysis was carried out using the Mann-Whitney U test. ***, P < 0.001; n.s., no significant difference. Dashed lines mark the detection limit.

In contrast, under liquid mating conditions with vigorous shaking (**Figure 4B**), the type IV pilus derivative displayed a marked improvement in transfer efficiency. Specifically, we observed a ∼10,000-fold increase in plasmid transfer from *E. coli* to *E. coli*, and an approximately tenfold increase in transfer to *Shigella*. In the case of *Salmonella*, conjugation remained below the detection limit for both plasmid versions under these conditions.

Together, these results show that incorporation of a type IV pilus can markedly enhance mating pair stabilization and conjugative transfer in liquid environments, provided that a compatible adhesin–recipient surface interaction is established between donor and recipient cells. By equipping stRP4 with the R64-derived type IV pilus, we converted a plasmid whose wild-type form conjugates inefficiently in liquid into a variant capable of robust transfer under these conditions, albeit in a recipient-dependent manner. This expands the functional repertoire of RP4-based tools by enabling efficient conjugation beyond solid surfaces, a property that may be particularly relevant in heterogenous environments such as the gut, where the relative contribution of solid-like versus liquid-like conditions remains unclear but where effective mating pair stabilization is likely to be a key determinant of plasmid dissemination^66^.

#### 3.3. Combining metabolic and transfer modules yields a multifunctional RP4 platform

Lastly, having demonstrated that the addition of the *fos* locus conferred a selective advantage under specific nutrient conditions and that the incorporation of the type IV pilus improved transfer efficiency in liquid environments, we generated a third plasmid carrying both traits (stRP4::*fos* locus + type IV pilus).

### 4. Conjugative delivery of the anti-pOXA-48 module reduces plasmid carriers in the population

Having developed and functionally validated several toxin–intein modules and engineered RP4 derivatives, we next integrated one of these systems into the conjugative platforms and assessed its activity against the resistance plasmid pOXA-48.

We first evaluated the system under a simplified and controlled *in vitro* setup, using 1:1 mating between two defined populations: (i) cells carrying an RP4 derivative (which act as RP4 donors and as recipients for pOXA-48) and (ii) cells carrying pOXA-48 (which serve as donors of this plasmid and as recipients of the RP4 variants). Because both the delivery platform and pOXA-48 are conjugative, we monitored the fate of both populations after reciprocal plasmid transfer (**Figure S10A**).

As shown in **Figure S10A**, introduction of the toxin–intein module resulted in a pronounced reduction in viability (4–5 log units) in both transconjugant populations, regardless of whether cells initially carried the delivery platform or pOXA-48. Importantly, this effect was not observed when the same experiments were performed using an isogenic strain lacking pOXA-48 (**Figure S10B**), demonstrating that killing strictly depends on the presence of the target plasmid. These results indicate that the specificity of the toxin–intein system is preserved after its integration into a fully conjugative platform, consistent with what was previously observed using mobilisable vectors.

While this setup allowed direct assessment of toxicity in defined transconjugant populations, we next asked how the toxin system affects the overall population in the absence of explicit selection for cells carrying both plasmids. This aspect is rarely addressed in conjugation studies, which typically focus exclusively on selected transconjugants.

To address this, we performed 24-hour solid-surface conjugation assays in *E. coli* populations carrying different RP4 derivatives, either with or without the anti-pOXA-48 module, co-incubated with an isogenic population harbouring pOXA-48. These experiments were conducted both in the presence and absence of scFOS in the medium, to assess the contribution of positive metabolic selection mediated by the *fos* locus in the corresponding plasmid variants.

As shown in **Figure S11**, the presence of the toxin module resulted in a measurable reduction in total colony-forming units even in the absence of selection for transconjugants (**Figure S11A**). Specifically, transfer of the anti-pOXA-48 module led to an approximately 1-log reduction in CFUs across all RP4 derivatives in the absence of scFOS (**Figure S11A, left panel**). When the plasmid carried the *fos* locus and scFOS was present in the medium, this reduction increased to up to ∼2 log units (**Figure S11A**, **right panel**). These results indicate that, even without explicit co-selection, conjugative delivery of the toxin–intein modules can significantly impact pOXA-48–carrying populations, and that positive metabolic selection further enhances this effect.

### 4. Positive metabolic selection enables conjugative toxin systems to block pOXA-48 invasion in complex communities

To better approximate the complexity expected in real applications, we next evaluated the performance of the engineered conjugative platforms in complex populations. Specifically, we used the OMM^12^ community^70^, a defined consortium of twelve bacterial species, which we supplemented with *E. coli* Mt1B1^70^ to increase diversity and to provide a potential recipient for plasmid transfer. As these experiments were conducted in liquid medium, we focused exclusively on the stRP4 derivatives encoding the type IV pilus, as these had previously been shown to display improved conjugation under liquid conditions (**Figure 4**).

Over the course of the experiment, *E. coli* populations were monitored at each time point by quantifying: (i) the total *E. coli* population (**Figure 5A**), (ii) cells carrying the stRP4 derivatives (**Figure 5B**), (iii) cells carrying pOXA-48 (**Figure 5C**), and (iv) cells co-carrying both plasmids (**Figure 5D**).

**Figure 5.**
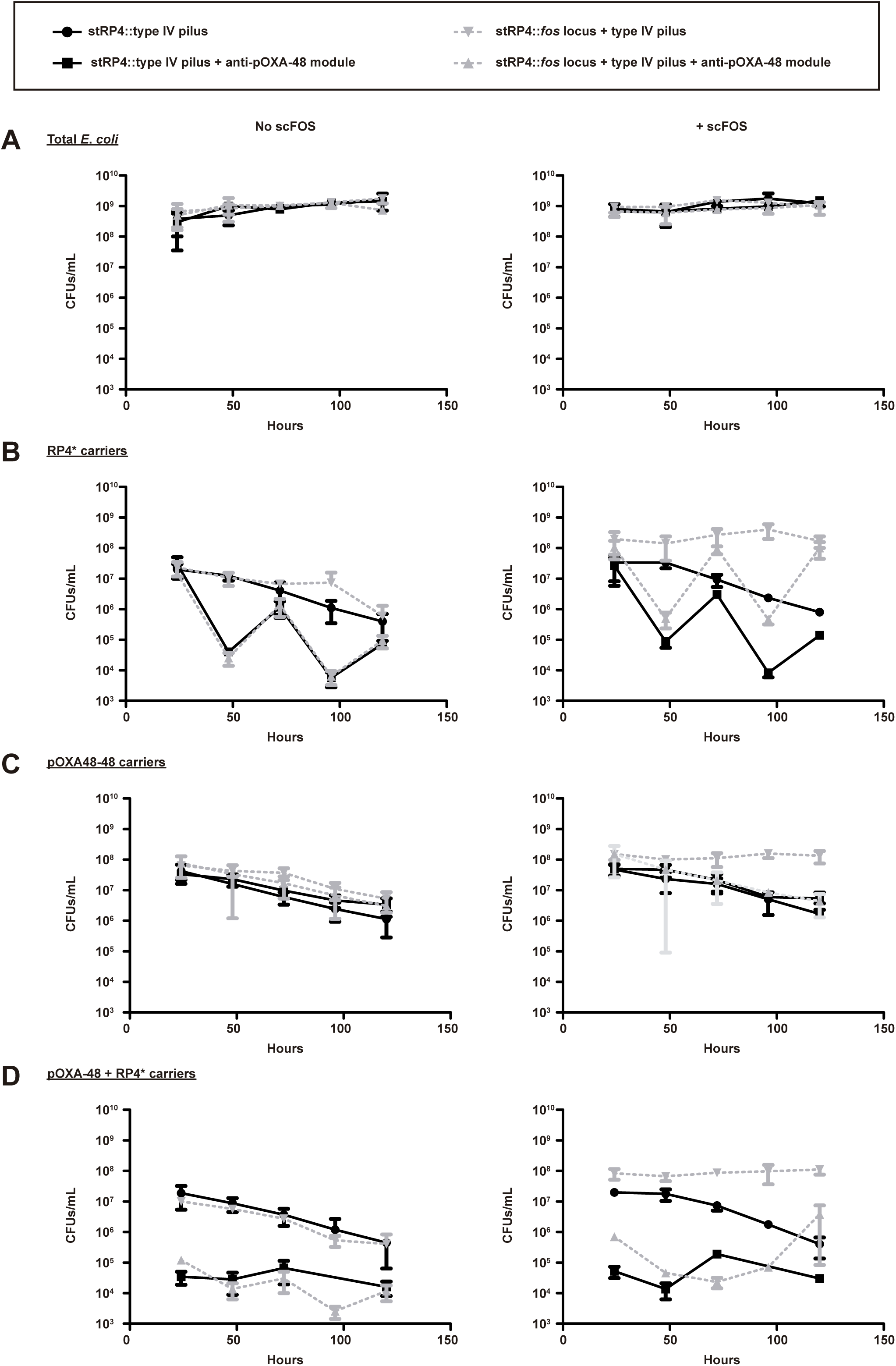
*E. coli* population dynamics in model communities during conjugative delivery of stRP4 derivatives. *E. coli* populations were monitored every 24 h over a five-day period in an OMM^12^-based community supplemented with *E. coli* Mt1B1. Conjugation assays were performed in liquid medium using *E. coli* MG1655 populations initially carrying either stRP4 derivatives or pOXA-48. stRP4 derivatives either carried or lacked the anti-pOXA-48 toxin–intein module. Panels are organised into two columns: experiments performed without scFOS (left) and with scFOS supplementation (right). Panel **A** shows total *E. coli* counts on non-selective medium. Panel **B** displays *E. coli* populations carrying stRP4 derivatives (kanamycin selection). Panel **C** shows populations carrying the pOXA-48 plasmid (carbenicillin selection). Panel **D** represents *E. coli* cells co-carrying both stRP4 derivatives and pOXA-48 (kanamycin and carbenicillin selection). Solid lines correspond to stRP4 derivatives encoding only the type IV pilus, whereas dashed lines represent derivatives additionally carrying the *fos* locus. Limit of detection: 10^2^ CFUs/ml

Across all conditions tested, the total *E. coli* population remained stable over time, showing no major fluctuations (**Figure 5A**).

In the absence of the toxin–intein module, both stRP4 and pOXA-48 carriers declined progressively over time (**Figure 5B** and **C**), despite stable total *E. coli* counts (**Figure 5A**). This pattern is consistent with *E. coli* Mt1B1, a strain known for its competitive performance within the OMM^12^ community^70^, being a poor recipient for conjugation under liquid conditions (**Supplementary Information 5**). This largely confines plasmid carriage to MG1655-derived populations, which appear to be progressively outcompeted by Mt1B1. In contrast, this decline was not observed for the stRP4 derivative carrying the *fos* locus under scFOS supplementation, where plasmid-bearing populations remained stable (**Figure 5B** and **C**, **right panels**).

When the toxin–intein module was present, pronounced fluctuations were observed specifically in the population carrying stRP4 derivatives (**Figure 5B**). This behaviour is consistent with an asymmetry in transfer dynamics, whereby the smaller pOXA-48 plasmid is transferred more rapidly into RP4-bearing cells than RP4 derivatives into pOXA-48 carriers, leading to preferential intoxication and loss of RP4-bearing cells. In line with this, no comparable fluctuations were observed in the pOXA-48–carrying population. Importantly, these fluctuations were attenuated when the stRP4 derivative carried the *fos* locus and scFOS was supplied (**Figure 5B**, **right panel**).

Regarding pOXA-48, in both the absence and presence of scFOS, stRP4 derivatives encoding only the type IV pilus failed to reduce the pOXA-48–carrying population at the population level (**Figure 5C**), unless cells co-carrying both plasmids were selectively monitored (**Figure 5D**). In contrast, the stRP4 derivative combining the *fos* locus and the type IV pilus displayed a distinct behaviour in the presence of scFOS. Under these conditions, and in the absence of the toxin–intein module, both plasmids were stably maintained over time, consistent with sustained co-carriage (**Figure 5C**, **right panel**). By contrast, when the anti-pOXA-48 module was present, the pOXA-48 population decreased by approximately 1.5 log units relative to the corresponding control, reaching levels comparable to those observed in the absence of scFOS or when using the type IV pilus–only derivative (**Figure 5C**, **right panel**). This pattern is consistent with the presence of a subpopulation refractory to pOXA-48 acquisition, likely cells already carrying the stRP4::fos–type IV pilus, which would be eliminated upon acquisition of the target plasmid. This effect was not observed with the type IV pilus–only derivative, likely because both plasmid-bearing populations declined over time under those conditions.

One additional observation worth noting is that, at later time points, particularly for the stRP4 carrying the *fos* locus and the type IV pilus in the presence of scFOS, an escaper population emerged and progressively increased under co-selection conditions (**Figure 5D**, **right panel**). This phenomenon mirrors what we observed in simpler *in vitro* systems (**Figures 1C** and **S5**) and highlights an important limitation that should be addressed in future designs.

Summing up, these experiments show that the performance of conjugative toxin–intein systems in complex microbial communities is shaped by plasmid transfer dynamics and the community context, including the availability of alternative recipients and metabolic exclusion of incoming strains. In the absence of co-selection for both plasmids, toxin delivery alone had limited impact on the long-term persistence of pOXA-48. By contrast, the inclusion of positive metabolic traits enabled sustained maintenance of cells carrying the delivery platform and, in combination with toxin activity, restricted pOXA-48 to its original carriers. Under these conditions, cells initially carrying pOXA-48 were progressively displaced by the resident microbiota, consistent with a competitive environment in which plasmids and hosts compete for available niches. At the same time, the emergence of escaper populations and the strong dependence on ecological conditions highlight key challenges for the deployment of conjugative antimicrobials in complex microbiomes, which we address in the next section.

## Discussion

This study builds on the work previously presented in López-Igual *et al*.^22^ and demonstrates that toxin-intein-based systems are an effective strategy for selective bacterial inactivation, provided that the biology of the target system is well understood. In-depth characterisation of species or element-specific regulatory networks enables the selection of highly selective promoter systems, ensuring that toxic activity is restricted to the intended context while minimising off-target effects. Once these key regulatory elements are identified, toxin-intein modules can be readily adapted to new targets. Furthermore, as these systems are orthogonal, they allow for modular and flexible designs^43,45^.

More precisely, in this work, we engineered toxin–intein cassettes targeting major enteric pathogens, including *Salmonella enterica* and *Shigella* spp., as well as the clinically relevant resistance plasmid pOXA-48. These systems were validated *in vitro* using standard conjugation assays and, in the case of pOXA-48, further evaluated in complex microbial communities, confirming their functionality across increasing levels of ecological complexity.

Our results demonstrate that the mode of antibacterial action, together with the biological nature of the target, shapes both efficacy and evolutionary outcomes, indicating that no single strategy is universally optimal. In *Shigella*, delivery of the bactericidal toxin CcdB imposed strong selective pressure. However, because the virulence plasmid pINV is highly enriched in insertion sequences, most escapers recovered carried deletions or rearrangements that rendered them avirulent. Thus, even when resistance emerged, it frequently came at the cost of pathogenicity, effectively neutralising the clinical relevance of escape events. This contrasts with the anti-pOXA-48 system, where escapers more commonly arose through mutations affecting the toxin–intein module itself, reflecting fundamental differences in the genetic architecture and evolutionary plasticity of the targeted element.

In *Salmonella enterica*, where we deliberately employed a bacteriostatic rather than bactericidal strategy, we demonstrate that virulence blockade can be as effective as cell killing in suppressing pathogenic potential. By interfering with the regulatory cascade required for T3SS expression, the system prevented the emergence of virulent subpopulations without inducing cell death. This inhibition-based strategy parallels CRISPRi approaches and is therefore expected to impose weaker selective pressure, reducing escaper frequencies while maintaining effective suppression of virulence-associated phenotypes.

In the case of the anti-pOXA-48 system, we validated the functionality and specificity of the toxin–intein module in clinical *Escherichia coli* and *Klebsiella* isolates, proving that its activity was strictly dependent on the presence of pOXA-48. In addition, we uncovered regulatory features that provide new insight into the biology of this clinically important plasmid. Our data suggest a previously unrecognised antagonistic interplay between the plasmid-encoded LysR_pOXA-48_ and the chromosomally encoded LysR_chr_, which may contribute to the adaptive behaviour of pOXA-48 across different host backgrounds.

In parallel to the development of these toxin-intein modules, we compared different plasmid-based conjugative strategies for their delivery. Each configuration with its inherent advantages and trade-offs. Mobilizable plasmids (setups 1 and 2) offer ease of engineering due to their smaller size but rely on the presence of a compatible conjugative element for transfer, which constrains their dissemination potential. In setup 2, depending on the order of transfer, mobilizable vectors may undergo multiple rounds of dissemination or be blocked by entry exclusion systems. In contrast, conjugative plasmids (setup 3) enable autonomous transfer and stable maintenance, as they encode complete transfer machineries, partitioning systems and post-segregational killing modules.

These differences in dissemination potential likely explain the behaviour observed in our experimental assays, in which conjugative plasmids consistently outperformed mobilisable vectors. To overcome the practical limitations associated with engineering large conjugative elements, we implemented a yeast-based assembly strategy that enabled efficient and flexible editing. This approach allowed the generation of a suite of engineered RP4 derivatives, including a streamlined backbone devoid of insertion sequences and transposons and with a simplified antibiotic resistance profile, and a variant capable of efficient conjugation in liquid environments. Together, these features expand the applicability of RP4-based delivery platforms beyond solid-surface assays and towards gut-relevant contexts.

An unexpected and particularly relevant finding was the impact of positive metabolic selection mediated by the *fos* locus. In the presence of its cognate substrate, scFOS, the *fos* locus not only enhanced host fitness but also stabilised plasmid-bearing populations within highly competitive ecosystems. This effect was evident even in complex communities, such as OMM^12^ supplemented with *E. coli* Mt1B1, a strain known for its strong competitive performance within this consortium^70^. Moreover, positive selection partially buffered both the fitness costs associated with plasmid carriage and the toxic effects linked to antimicrobial module deployment, underscoring the importance of coupling the delivery of a given trait to ecological competitiveness.

When integrating the anti-pOXA-48 toxin–intein module into conjugative delivery platforms, in simplified solid-surface mating assays, we observed a pronounced effect on the population of pOXA-48 carriers even in the absence of explicit selection for transconjugants. Such effects are rarely captured in classical conjugation studies that focus exclusively on transconjugants. This phenomenon was less evident in liquid mating assays within the OMM^12^ consortium, likely reflecting differences in cell density and mating dynamics that enable extensive reciprocal plasmid exchange.

When evaluated in complex communities under liquid conditions, the system primarily limited the dissemination of pOXA-48 rather than achieving its complete elimination. This behaviour resembles naturally occurring CRISPR-based defence systems that prevent the establishment of incoming mobile genetic elements without necessarily clearing them from the environment. These observations suggest that conjugative toxin–intein systems can function as plasmid traps: when the delivery platform is present in the microbiota, by coupling toxin activation to plasmid entry, they prevent incoming plasmids from establishing within already adapted members of the resident microbiota, thereby blocking population-level invasion.

An additional factor likely contributing to this behaviour is the asymmetry in transfer kinetics between the two conjugative elements. The smaller pOXA-48 plasmid may be transferred more rapidly into RP4-bearing cells than RP4 derivatives are transferred into pOXA-48 carriers, leading to preferential intoxication of delivery-platform hosts. This asymmetry provides a plausible mechanistic explanation for the pronounced fluctuations observed in RP4-carrying populations and highlights the importance of considering reciprocal transfer dynamics when designing conjugative interference strategies.

Based on this, beyond toxin-based approaches, conjugative platforms could also be repurposed to interfere with plasmid dissemination through exclusion rather than killing. Engineering delivery plasmids to encode batteries of entry or surface exclusion systems could prevent the uptake of specific incompatibility groups without imposing direct lethality. In this configuration, such systems would act as effective barriers against the acquisition of new resistance elements by selectively impairing the dissemination of incoming plasmids. Importantly, by breaking the symmetry of reciprocal transfer, exclusion-based strategies would favour the mobility of the delivery platform while restricting the spread of the target plasmid, thereby complementing toxin- and CRISPR-based approaches. This mode of action would primarily affect newly arriving donors, limiting plasmid invasion without imposing strong selective pressure on the resident microbiota carrying the delivery platform.

Despite these advances, there are several areas in which further optimization is possible. One promising direction is the incorporation of anti-defence systems into stRP4^71^, which could help overcome recipient immunity barriers, enhancing conjugation efficiency and broadening host range. Targeting specificity could also be enhanced using nanobodies^72^ or by manipulating the shufflon system, either by integrating pre-existing cassettes or through engineering of the pilus tip using approaches such as random mutagenesis^73^ or <u>d</u>iversity-<u>g</u>enerating retroelements (DGRs)^74^.

Another way to strengthen efficacy would be to minimize the emergence of escapers by combining multiple antibacterial systems. A recent study demonstrated that the strongest suppression of bacterial growth occurred when CRISPR-Cas9 was paired with one or two toxin genes^75^. Along these lines, modules integrating multiple antibacterial systems with distinct mechanisms of action -such as toxins, CRISPRi^49^ or CRISPR-associated transposases^76^-could provide a more robust barrier against the development of resistance.

Manipulating plasmid copy number could also represent an attractive, yet challenging, strategy to boost toxin activity. Increasing copy number specifically within the target organism could enhance the effect of the toxin through higher gene dosage. This could be achieved either by engineering copy number control circuits^77^ or by exploiting natural regulatory mechanisms^78^. Alternatively, disabling plasmid entry exclusion systems would allow repeated plasmid uptake and transient increases in copy number. Such repeated delivery could also be advantageous in scenarios where mutations in the toxin render the resident plasmid ineffective, as incoming functional copies could restore activity. This strategy, however, is not without important drawbacks: the absence of exclusion mechanisms can lead to lethal zygosis^79,80^ and transient copy number surges may increase off-target activity. Careful evaluation would therefore be required before implementing these strategies.

In the longer term, engineering stRP4 as a shuttle vector capable of replicating and transferring across additional gut-associated bacterial clades, such as *Bacteroides* or *Firmicutes*, could substantially broaden the applicability of the system. Stable maintenance in these taxa could be achieved by integrating replicons and partition systems from native plasmids, while providing *oriT* and relaxase modules from successful parasites might enable efficient secondary dissemination^81^.

A major consideration when deploying synthetic plasmids is the potential risk of uncontrolled spread within microbial communities. To mitigate this, multiple biocontainment layers can be envisioned. One option involves the use of T4SS-dependent bacteriophages to selectively eliminate plasmid-bearing cells after therapeutic use^82^. Alternatively, conditional toxin activation via inteins or environmentally responsive promoters could restrict activity to specific niches. For example, placing the expression of an antitoxin under a temperature-dependent promoter, such as that of *virF*, would inhibit the action of a constitutively produced toxin in host-associated settings but unleash it upon environmental exposure. In parallel to plasmid containment strategies, the use of probiotic strains as donors could further reduce the risk of dysbiosis by occupying the niche left by the eliminated pathogen.

Taken together, our results represent a step forward in the development of highly specific antimicrobial strategies based on conjugative systems^83^. By combining targeted toxin–intein modules with engineered delivery platforms that explicitly account for ecological competitiveness and transfer dynamics, we establish programmable systems capable of selectively interfering with pathogen and resistance dissemination at the community level. While our study focused on *in vitro* and model-community settings, a logical next step will be to evaluate these strategies *in vivo*, where additional layers of complexity -including host-associated factors, spatial structure and microbiota heterogeneity-are expected to further shape conjugation outcomes and define the limits of these systems.

## Material and methods

### Bacterial strains, plasmids and oligonucleotides

Bacterial strains, plasmids, and oligonucleotides used in this work are listed in Tables S1, S2, and S3, respectively.

### Culture conditions

*Escherichia coli* and *Salmonella* enterica strains were routinely grown in LB Lennox at 37°C unless otherwise indicated. *Shigella* strains were routinely grown in LB Lennox at 30°C unless otherwise indicated. When required, media were supplemented with the appropriate reagents at the following concentrations: carbenicillin (Carb), 100 μg ml^−1^; chloramphenicol (Cm), 25 μg ml^−1^; kanamycin (Kn), 50 μg ml^−1^; apramycin (Apra), 30 μg ml^−1^; Tetracycline (Tet), 15 μg ml^−1^; ampicillin (Amp), 100 μg ml^−1^; diaminopimelic acid (DAP), 0.3 mM; scFOS, 0.5% (w/v); arabinose, 0.2% (w/v); glucose, 1% (w/v). Bacteriological agar was used as a gelling agent.

### DNA manipulations

Routine DNA manipulations were performed using standard procedures. Oligonucleotides were synthesized by Eurofins (Cologne, Germany). All constructed plasmids were confirmed by Nanopore sequencing.

### Construction of RP4 derivatives

All RP4 derivatives, with the exception of the AmpR variants that were generated by λ-Red recombination to replace the *aphA* gene with *bla*, were constructed by yeast-mediated assembly following the protocol described in Ibarra-Chávez *et al*.^84^. After assembly, extraction, and transfer to *E. coli*, the <u>y</u>east <u>a</u>rtificial <u>c</u>hromosome (YAC) was removed as follows. The plasmid pBAD33::ΦC31Int was introduced into *E. coli* carrying the corresponding RP4 derivative. Cultures were then grown to exponential phase, induced with arabinose for 3–4 hours, and subsequently plated on Kn-containing medium. Resulting colonies were streaked on Kn and Kn + Carb plates to screen for YAC excision.

### Stability assays

Plasmid stability was evaluated in *E. coli* MG1655 carrying either RP4 or stRP4 over a 14-day period. Cultures were serially diluted 1:1000 into fresh LB Lennox medium every 12 hours and incubated at 37 °C with shaking. Every 24 hours, aliquots were plated in parallel on LB Lennox and on LB Lennox-Kn plates. The proportion of plasmid-bearing cells was estimated by comparing colony counts on selective versus non-selective plates. For additional confirmation, 100 colonies from non-selective plates were replica-plated on both media. Experiments were performed in triplicate for each plasmid variant.

### Conjugation assays on solid media

Donor and recipient strains were grown to an OD_600_ of 0.5 and mixed at a 1:1 ratio. A 50 µL aliquot of the mixture was spotted onto a nitrocellulose filter placed on an LB Lennox–DAP agar plate and allowed to dry. The conjugation mixture was incubated for 3 h at 37 °C. Following incubation, the biomass was resuspended in PBS (1x), serially diluted and plated onto selective media as follows: LB Lennox agar for enumeration of total potential recipients, LB Lennox–DAP–Apra for donor counts, and LB Lennox supplemented with the corresponding antibiotic marker of the plasmid under study (Kn, Carb and/or Cm) for quantification of transconjugants. Transfer frequencies were calculated as transconjugants per recipient cells.

### Growth assays in scFOS

Overnight cultures were prepared in Mueller-Hinton (MH) supplemented with Kn. The following day, cells were harvested, washed three times with sterile fresh medium and adjusted to an OD_600_ of 1. Serial dilutions were prepared in MH-Kn, and 200 µL of the 1:1000 dilution were put to grow. Cultures were incubated at 37 °C for 48h under shaking conditions, and bacterial growth was monitored.

### Competition assays in the presence or absence of scFOS

Overnight cultures were grown in MH. The following day, cultures were diluted 1:1000 into 1.5 ml of fresh medium in 24-well plates and propagated for 5 consecutive days, with dilutions performed every 12 h. To monitor the dynamics of the competing strains, aliquots were taken every 24 h and plated onto selective media containing either Kn or Carb, allowing quantification of each plasmid-bearing population. All incubations were carried out at 37 °C under shaking conditions.

### Liquid conjugation assays

Donor and recipient strains were grown in LB Lennox medium to an OD_600_ of 0.5 and subsequently diluted 1:1000 into 50 ml of fresh medium. Conjugation mixtures were incubated for 3 h at 37°C with shaking. Following incubation, cultures were serially diluted in PBS and plated onto selective media to determine the different bacterial populations: LB Lennox for total recipients, LB Lennox–DAP–Apra for donors, and LB Lennox-Kn for transconjugants. Transfer frequencies were calculated as transconjugants per recipient cells.

### Reporter validation in *Shigella* spp

Overnight cultures of *Shigella* spp. and *E. coli* MG1655, both carrying either the empty or reporter plasmid, were grown in LB Lennox-Kn at 30°C with shaking. The following day, 20 mL of fresh selective medium were inoculated at a 1:100 dilution. Cultures were incubated for 3 h either at 30°C or at 37°C, both under shaking conditions. After incubation, fluorescence levels were measured by FACS.

### Reporter validation in *Salmonella enterica*

Overnight cultures of *Salmonella enterica* strains and *E. coli* MG1655, both carrying either the reporter plasmids or the empty vector, were grown in LB Miller (0.17 M NaCl) supplemented with Kn at 37°C with shaking. The following day, 5 mL of fresh medium consisting of LB Miller supplemented with 0.08 M NaCl, 10 mM MgSO_4_ and Kn were inoculated at a 1:100 dilution. Cultures were incubated at 37°C with shaking. Samples were collected at 4h, 8h and overnight. Fluorescence was quantified by FACS.

### Reporter validation for pOXA-48

Two versions of the *P_pfp_*_-*ifp*_-*gfp* reporter plasmid were built: one with an intact copy of *lysR_chr_*, mimicking the exact chromosomal locus of the *pfp*-*ifp* operon, and a second one with a methionine-less version of *lysR_chr_*. Overnight cultures of *E. coli* MG1655, *K. pneumoniae* KPN15 and *C. freundii* CF13, carrying either version of the reporter plasmid and carrying / not-carrying pOXA-48 were grown in LB Lennox at 37°C with shaking. The following day, fresh cultures were inoculated at a 1:100 dilution and incubated for 3 h at 37°C under shaking conditions. After incubation, fluorescence levels were measured by FACS.

### Killing assays in *Shigella*

Overnight cultures of *Shigella* spp. (recipients) and *E. coli* NGE*pir* carrying either the empty vector or the plasmid encoding the anti-Shigella toxin-intein system (donors) were grown at 30°C under shaking conditions. The following day, cultures were diluted 1:100 in fresh medium and incubated at 30°C with shaking until reaching an OD_600_ of ∼0.5. Donor and recipient cultures were collected, washed, mixed at a 1:1 ratio and 50 µL of the mixture was spotted onto a nitrocellulose filter placed on LB Lennox agar supplemented with DAP. After drying, plates were incubated for 3 h at either 30 °C or 37 °C. Following incubation, the biomass was recovered by resuspension in PBS, serially diluted, and plated onto selective media: LB Lennox (for total recipients), LB Lennox-DAP-Apra (for donors), and LB Lennox-Kn (for transconjugants). Plates were incubated at both 30 °C and 37 °C before enumeration.

### Blocking virulence assays in *Salmonella*

Overnight cultures were prepared in 5 mL of LB Miller supplemented with: Carb for *Salmonella* strains harbouring pSC101-reporter plasmid or Kn-DAP for *E. coli* NGEpir strains carrying either the empty vector or plasmid with the anti-Salmonella toxin–intein module. Cultures were incubated at 37 °C with shaking. The following day, overnight cultures were washed and the OD_600_ adjusted to 0.5. Donor and recipient cells were mixed in a 1:1 ratio and 50 µL of the mixture were spotted onto a nitrocellulose filter placed on a LB Lennox-DAP plate. Drops were allowed to dry and plates were incubated for 2 h at 37 °C. After incubation, the biomass was recovered in pre-warmed (37 °C) LB Miller supplemented with 0.08 M NaCl, 10 mM MgSO_4_, Kn and Carb. Bacterial suspensions were incubated at 37 °C with shaking. For monitoring the fluorescence levels, culture samples were collected at 6, 7, 8, 9, 10 and 24h, and analyzed by FACS.

### Killing assays pOXA-48

*E. coli* NGEpir carrying pSEVA228::anti-pOXA-48 or pSEVA228 were used in the conjugation assays as donor strains. *E. coli* MG1655, and clinical strains of *E. coli* (C288), *K. pneumoniae* (K147) and *C. freundii* (CF13), all carrying pOXA-48 or not carrying the plasmid were used as recipient strains. Conjugation assays were performed as described for the killing assays in Shigella, with the difference that cultures were always incubated at 37°C.

### *In vitro* experiments in complex populations

First, the OMM^12^ community^70^ was grown anaerobically in Anaerobic Akkermansia Medium over four days^85^ to reach stable community composition. Experimental strains (MG1655 with stRP4::type IV pilus, stRP4::type IV pilus + anti-pOXA-48 module, stRP4::type IV pilus + *fos* locus, stRP4::type IV pilus + *fos* locus + anti-pOXA-48 module or pOXA-48 and *E. coli* Mt1B1^70^) were introduced after community passage with an adjusted OD 1.0 in a 1:100 ratio (10 µl culture in 1000 µl fresh Anaerobic Akkermansia Medium with or without 0.5% scFOS). In total four experimental groups were tested with three technical replicates (3 wells) each. The community with the experimental strains was passaged every 24 h in a 1:100 ratio into fresh medium. After every passage, grown community samples were exported, serially diluted in PBS and plated onto selective media to determine CFUs of the different bacterial populations (10 µl spots on MacConkey with Kn for stRP4 carriers; and Amp for pOXA-48 carriers).

### Statistics and reproducibility

All experimental data are representative of the results of at least three independent biological repeats. All replication attempts were successful. Statistical analysis of experimental data was done using GraphPad Prism 9.1.2 (GraphPad Software, Inc., CA, USA).

## Data availability

The data, plasmids and strains generated for this study are available upon request.

## Acknowledgements

P.D.-M. was supported by the European Commission through a Marie Skłodowska-Curie Postdoctoral Fellowship (HORIZON-MSCA-2022-PF-01 CoKiP, grant agreement 101104168). A.C.-V. was supported by an EMBO postdoctoral fellowship (ALTF 322-2022). M.L. was supported by the Fondation pour la Recherche Médicale FDM202106013531. Work in the A.S.M. lab was supported by MCIN/AEI/10.13039/501100011033 and the European Union NextGenerationEU/ PRTR (Project PCI2021-122062-2A) and by the European Research Council (ERC) under the European Union’s Horizon Europe research and innovation programme (ERC-2022-CoG Project 101086992–PLAS-FIGHTER). D.M.’s laboratory is funded by the Institut Pasteur and the Centre National de la Recherche Scientifique. This work was supported by the Fondation pour la Recherche Médicale Equipe FRM (EQU202103012569). This work was also supported by Pfizer Innovation France, the French Government’s Investissement d’Avenir program Laboratoire d’Excellence ‘Integrative Biology of Emerging Infectious Diseases’ [ANR-10-LABX-62-IBEID] and the project MOB-TARGET has been supported by ANR (ANR-21-AAMR-0006) under the framework of the JPIAMR - Joint Programming Initiative on Antimicrobial Resistance-(JPIAMR2021-09).

## Author contributions

Conceptualization: P.D.-M., D.M.

Methodology: P.D.-M., A.C.-V.

Investigation: P.D.-M., A.C.-V., B.A., L.N., M.L., M.D.-G.

Formal analysis: P.D.-M., A.C.-V., B.A., L.N.

Writing – original draft: P.D.-M., A.C.-V.

Writing – review & editing: all authors.

Supervision: P.D.-M, B.S., J.-M-G., Á.S.M., D.M.,

Funding acquisition: P.D.-M., B.S., Á.S.M., D.M.

All authors approved the final version of the manuscript.

## Competing interests

The authors declare no competing interests.

## Figure descriptions

**Figure S1.**
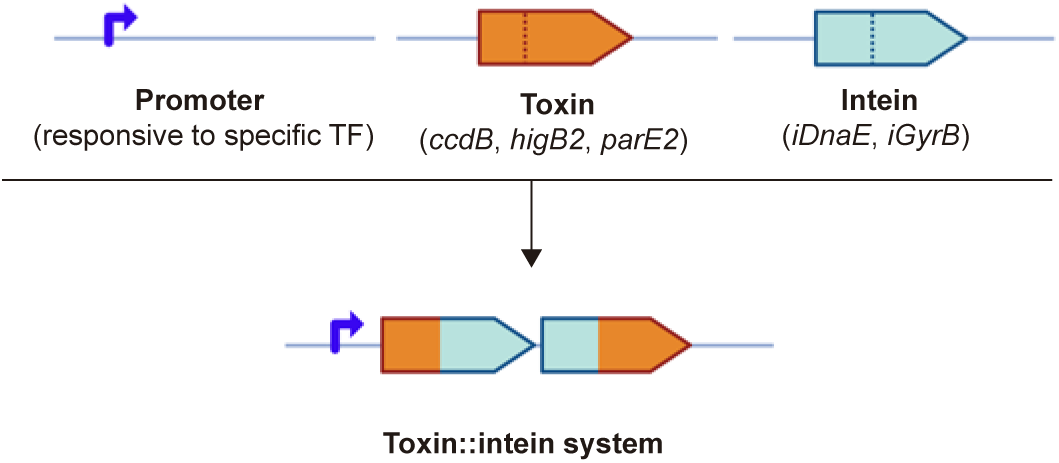
Schematic representation of the toxin–intein systems used in this study. Each system consists of a promoter responsive to specific transcription factors driving expression of a two-ORF operon. The first ORF encodes the N-terminal fragment of the toxin fused to the N-terminal fragment of the intein, whereas the second ORF encodes the C-terminal fragment of the intein fused to the C-terminal fragment of the toxin.

**Figure S2.**
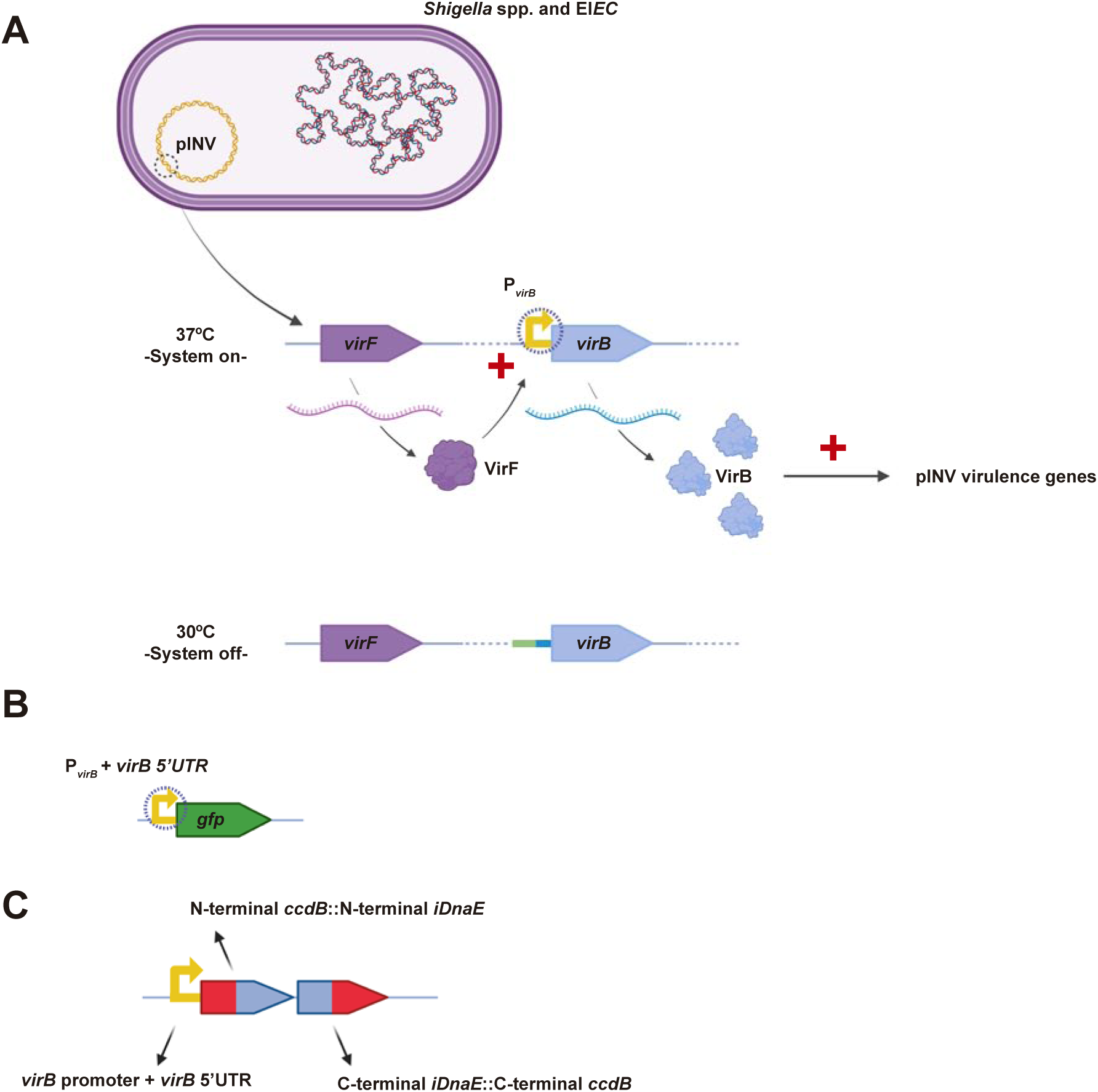
Schematic representation of (**A**) the *Shigella* regulatory mechanism exploited for the development of the anti-*Shigella* toxin–intein system; (**B**) the reporter used to assess the responsiveness of the *virB* promoter; and (**C**) the toxin–intein module generated for *Shigella* targeting.

**Figure S3.**
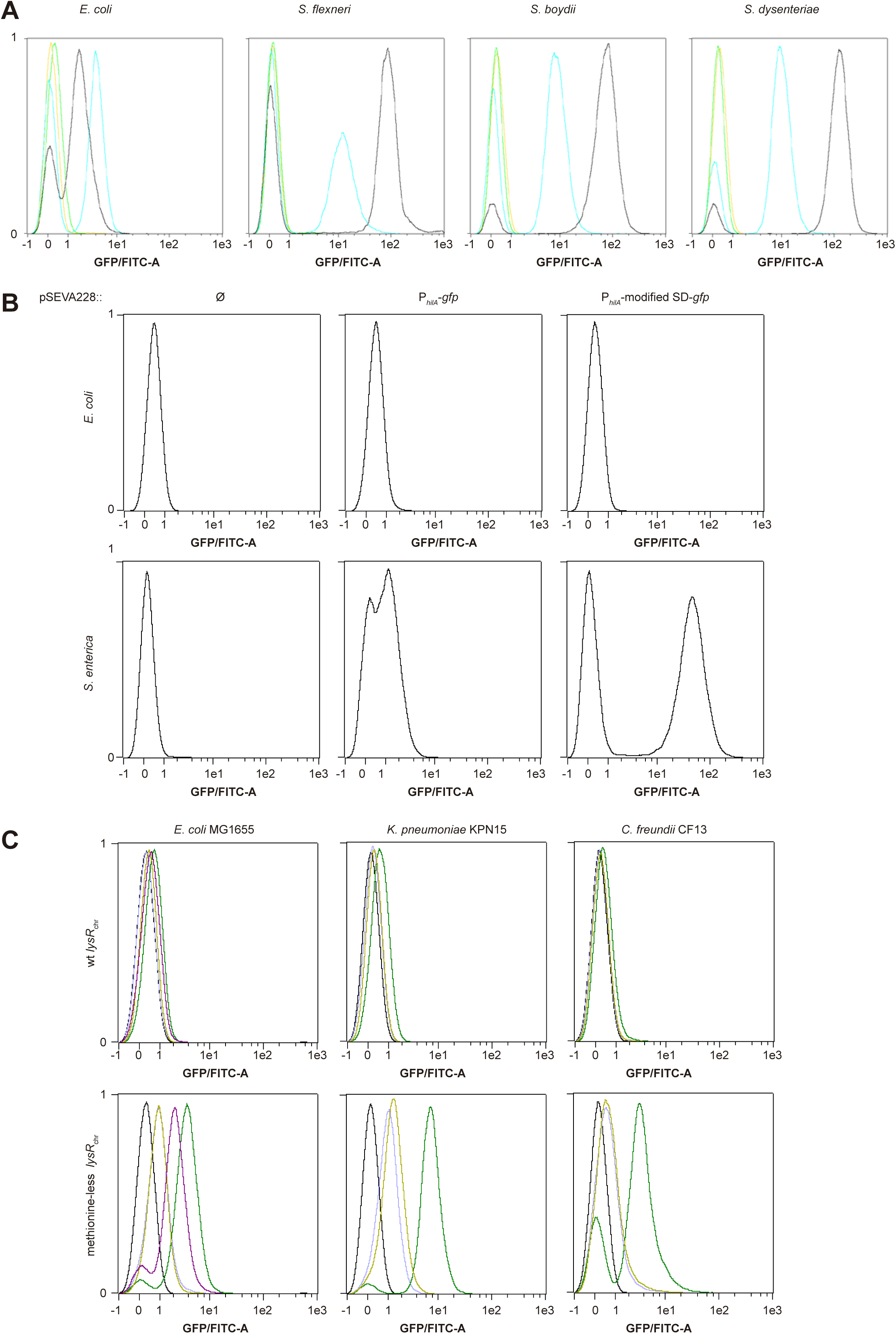
Evaluation of target-specific promoter responsiveness for *Shigella*, *Salmonella* and pOXA-48 using flow cytometry. **A.** Activity of the *virB* promoter in *E. coli* MG1655 (control) and three *Shigella* species (*S. flexneri*, *S. boydii* and *S. dysenteriae*), each transformed with either an empty plasmid or a plasmid carrying a P*_virB_*–*gfp* reporter. Fluorescence was measured at 30°C and 37°C to assess temperature-dependent promoter activation. **Color code**: yellow, empty plasmid at 30°C; light green, empty plasmid at 37°C; light blue, reporter plasmid at 30°C; black, reporter plasmid at 37°C. **B.** Evaluation of *hilA* promoter activity in *E. coli* MG1655 (top) and *Salmonella enterica* serovar Typhimurium SL1344 (bottom). Strains carrying: (left) empty vector, (middle) reporter with the native RBS or (right) reporter with an optimized RBS. Fluorescence was measured under *hilD*-inducing conditions. **C.** Evaluation of the *pfp*-*ifp* promoter’s responsiveness to pOXA-48 in *E. coli* MG1655, *K*. *pneumoniae* KPN15 and *C*. *freundii* CF13. Each strain transformed with a plasmid carrying a P*_pfp_*_-*ifp*-_*gfp* reporter, preceded by either a wild-type version of *lysR_chr_*, or a methionine-less version. Color code: purple, GFP positive control; black, GFP negative control; dark green, presence of pOXA-48; light green, presence of pOXA-48 Δ*lysR*; light blue, absence of pOXA-48. The data shown in this figure are representative of at least three independent biological replicates.

**Figure S4.**
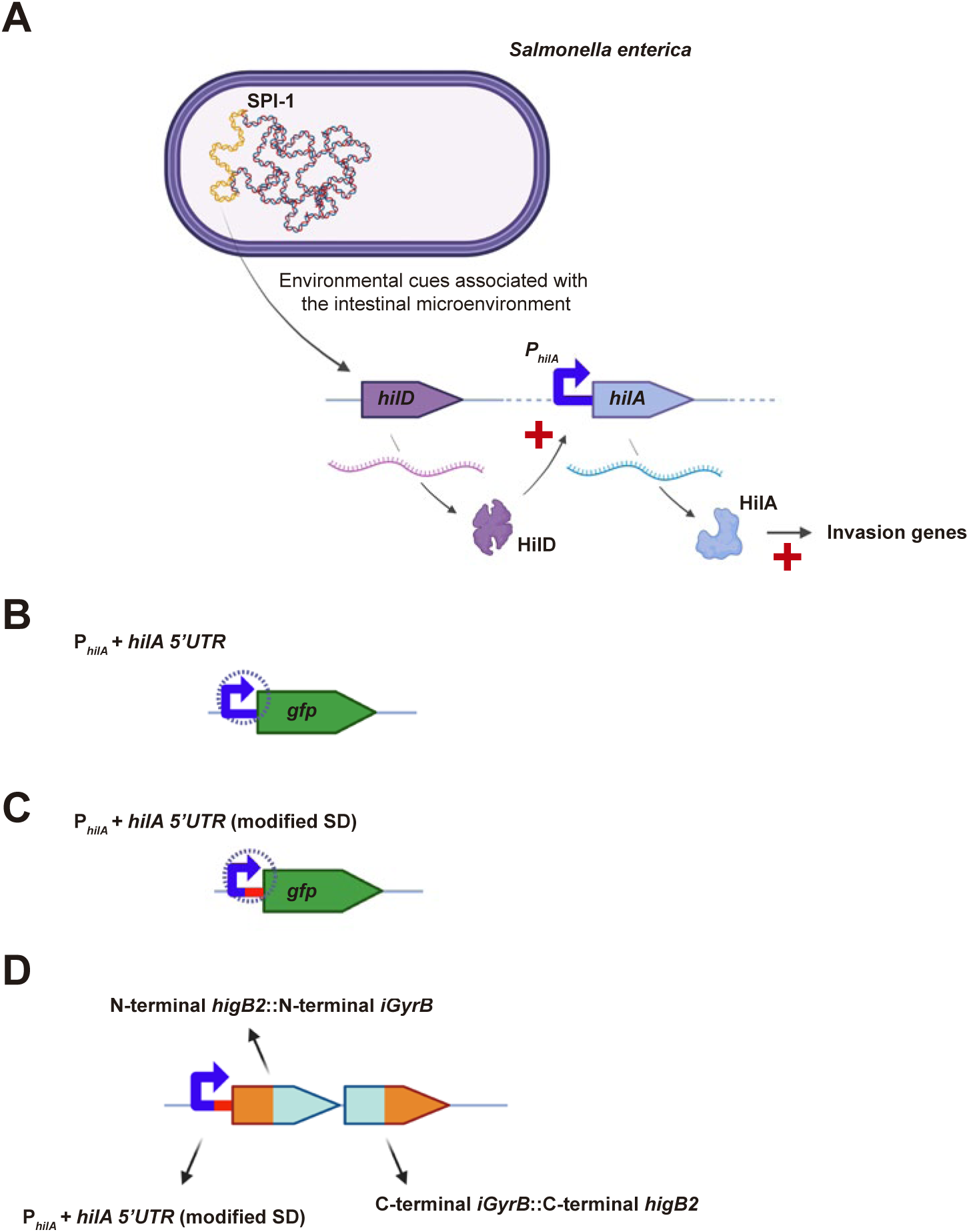
Schematic representation of (**A**) the *Salmonella enterica* regulatory mechanism exploited for the development of the anti-*Shigella* toxin–intein system; (**B,C**) the reporters used to assess the responsiveness of the *hilA* promoter; and (**D**) the toxin–intein module generated for *Salmonella* targeting.

**Figure S5.**
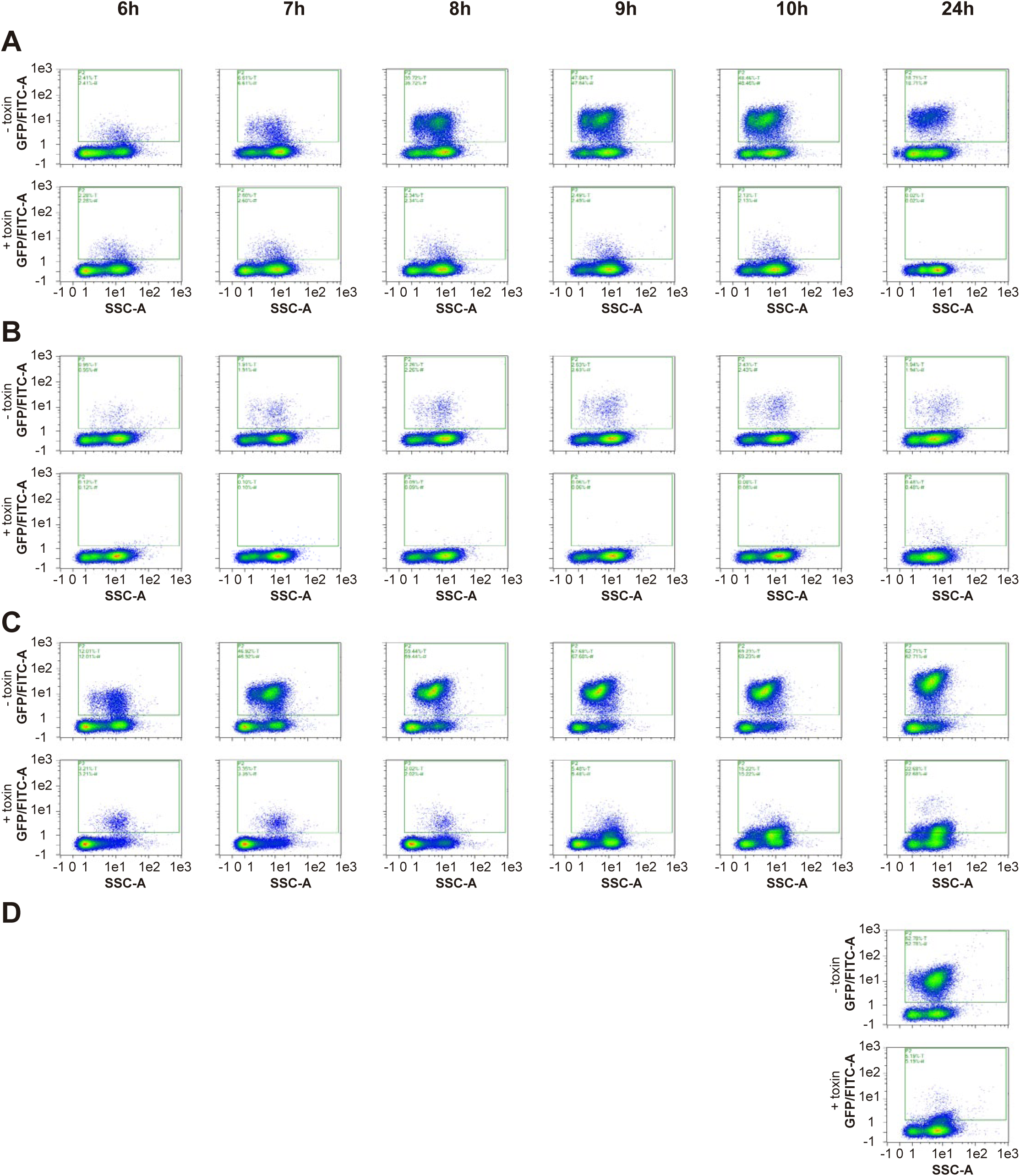
*hilA* promoter kinetics in different *Salmonella enterica* strains. Time-course analysis of *hilA* promoter activity at 6, 7, 8, 9, 10 and 24 h post-conjugation. Strains tested: **A.** Typhimurium, **B.** Typhi, **C.** Paratyphi A and **D.** Gallinarum. Recipient cells received either the empty vector (upper panels) or the plasmid carrying the toxin–intein module (lower panels). Reporter fluorescence in transconjugants was quantified by flow cytometry under *hilD*-inducing conditions. **Note**: for Gallinarum, only the 24-h time point is shown due to its slower growth. The data shown in this figure are representative of at least three independent biological replicates.

**Figure S6.**
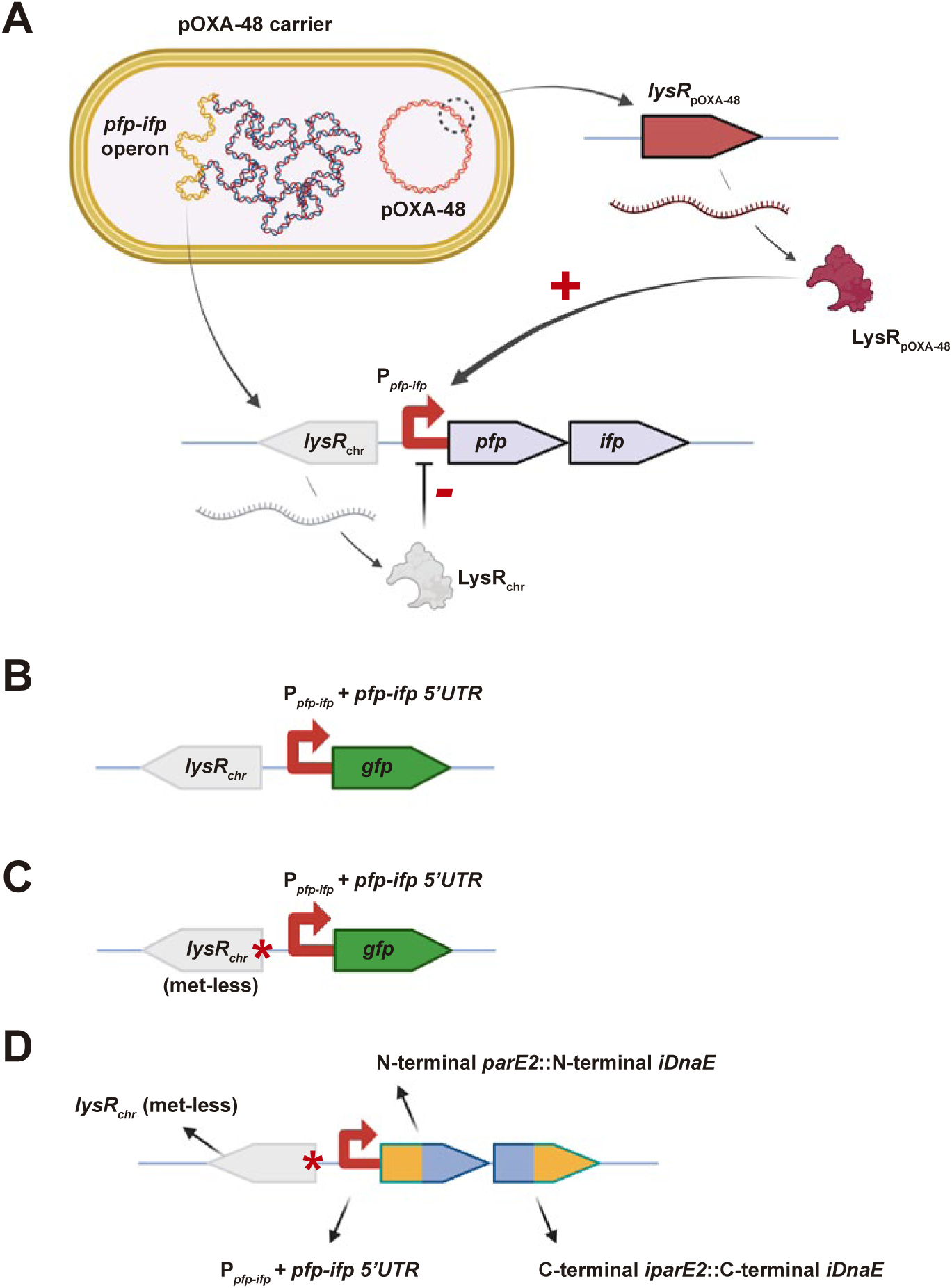
Schematic representation of (**A**) the pOXA-48 regulatory mechanism exploited for the development of the anti-pOXA-48 toxin–intein system; (**B,C**) the reporters used to assess the responsiveness of the *pfp-ifp* promoter; and (**D**) the toxin–intein module generated for targeting pOXA-48 carriers.

**Figure S7.**
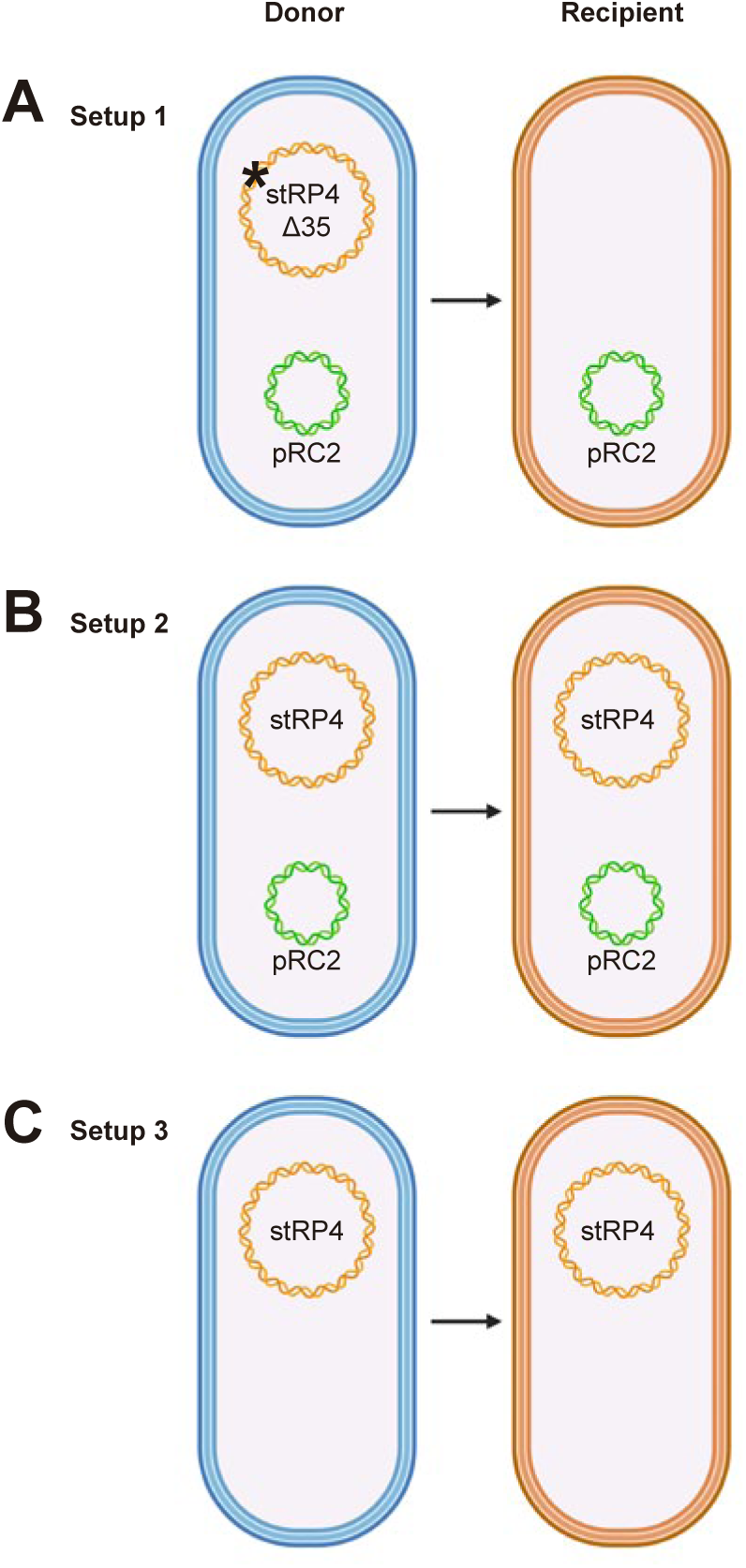
Schematic representation of the three plasmid configurations used to experimentally assess which setup achieves the highest conjugative transfer efficiency. For each configuration, the plasmid arrangement in the initial donor strain (**left**) and the expected outcome after conjugation, based on the transfer or mobilization capacity of each plasmid (**right**), are shown. **A**. Setup 1: the mobilizable plasmid pRC2 was used in combination with a streamlined RP4 derivative carrying a 15-nucleotide deletion in the *oriT* sequence (stRP4 Δ35), which abolishes RP4 self-transfer while preserving its ability to function as a conjugative helper. **B**. Setup 2: pRC2 was used together with the streamlined RP4 variant (stRP4), allowing both plasmids to be mobilized during conjugation. **C**. Setup 3: the streamlined RP4 derivative (stRP4) was used alone as a self-transmissible plasmid.

**Figure S8.**
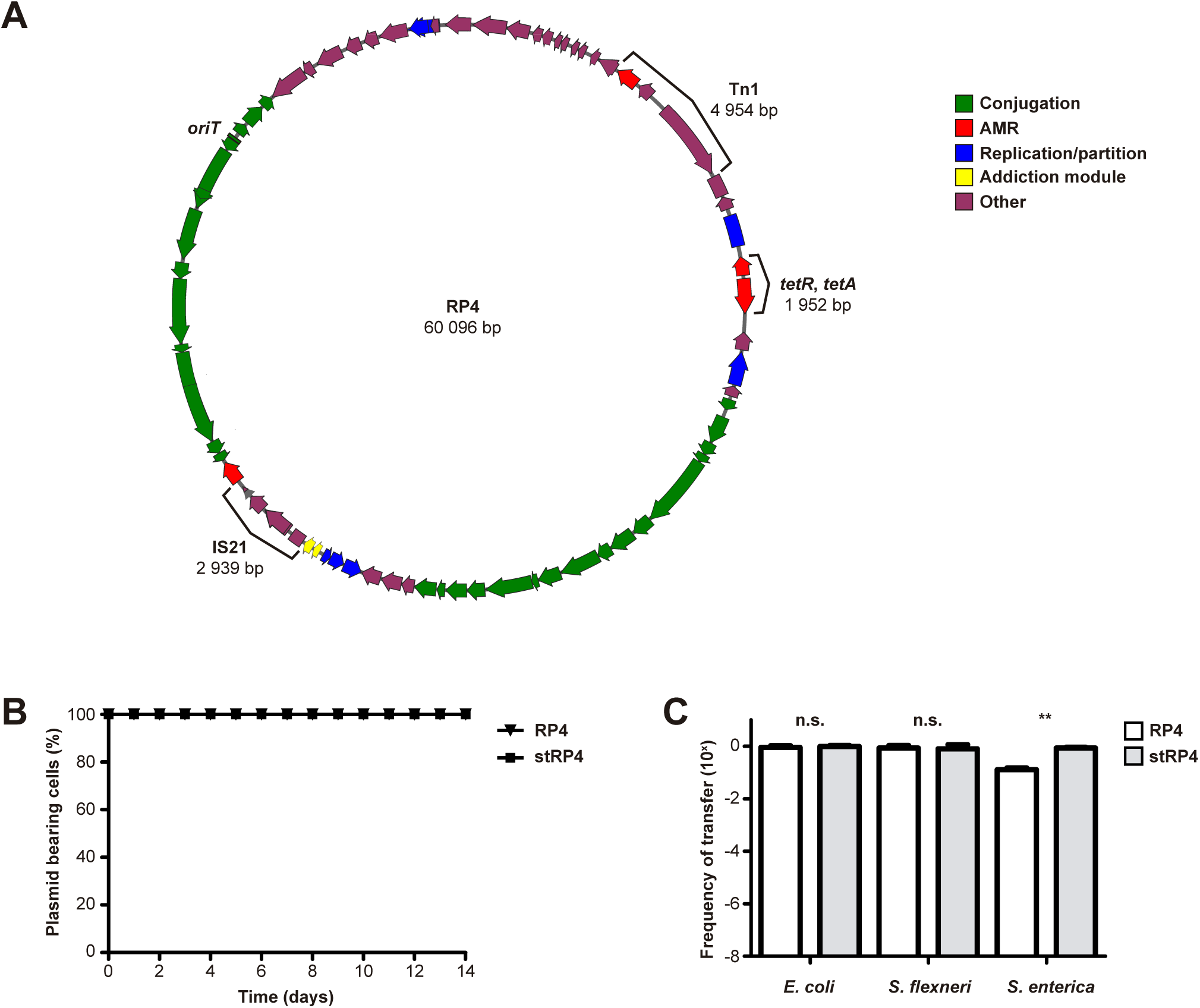
A. Schematic representation of the wild-type RP4 plasmid. Brackets indicate the regions that were removed in the RP4 streamlined version (stRP4). **B.** Plasmid stability of RP4 and stRP4 in *E. coli* MG1655 over a 14-day period. Each day, cultures were plated on LB agar and LB agar supplemented with kanamycin. Colony counts were used to estimate plasmid retention. 100 colonies from non-selective plates were replica-plated onto both selective and non-selective media to assess plasmid presence. The percentages shown correspond to the fraction of colonies that retained kanamycin resistance among the 100 tested. Data represent mean values and standard deviation from three independent biological replicates. **C.** Comparison of plasmid transfer efficiencies of RP4 and stRP4 using solid surface conjugation assays. An auxotrophic *E. coli* strain was used as donor and *E. coli* MG1655, *Shigella flexneri* 2a or *Salmonella enterica* SL1344 as recipients. Statistical analysis was carried out using the Mann-Whitney U test. ***, P < 0.001; n.s., no significant difference.

**Figure S9.**
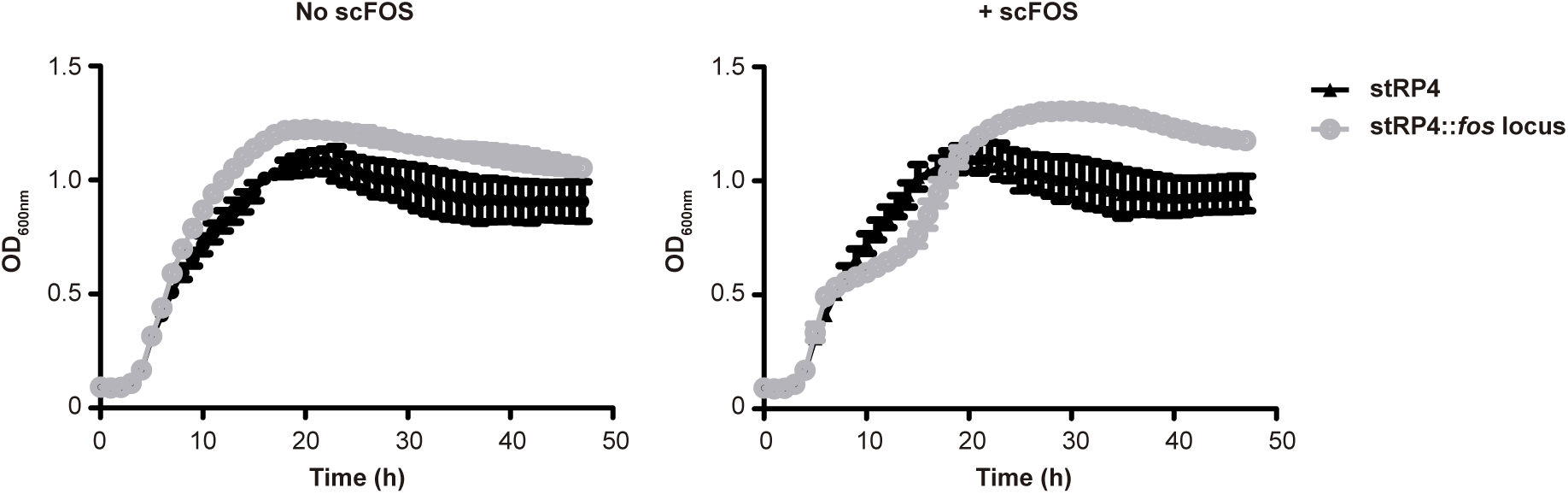
Effect of the *fos* locus on *E. coli* MG1655 growth. Growth curves of *E. coli* MG1655 carrying either the stRP4 plasmid (black) or its derivative containing the *fos* locus (grey), in the absence (left) and presence (right) of exogenously added scFOS (0.5% w/v). Data represent mean values from six independent biological replicates. Error bars indicate standard deviation.

**Figure S10.**
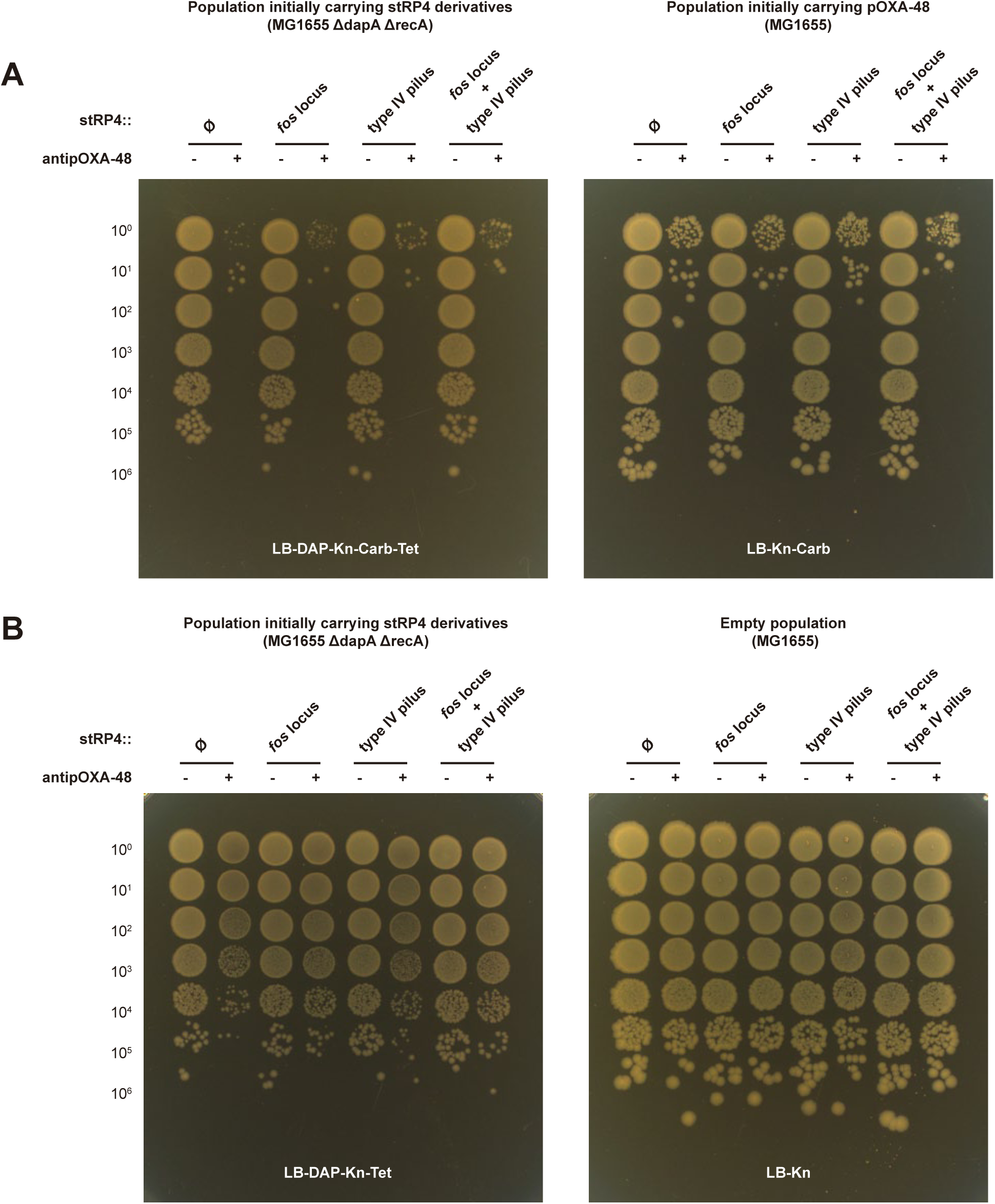
*In vitro* validation of the toxin–intein module against pOXA-48 when integrated into the final conjugative platforms. **A.** Because both the delivery plasmids and pOXA-48 are conjugative, we tracked the fate of two transconjugant populations after mating: cells initially carrying the delivery platform that subsequently acquired pOXA-48 (left panels), and cells initially carrying pOXA-48 that received the delivery platform (right panels). **B.** To verify that specificity was maintained, parallel experiments were performed with an isogenic *E. coli* MG1655 strain lacking pOXA-48. The data shown in this figure are representative of at least three independent biological replicates.

**Figure S11.**
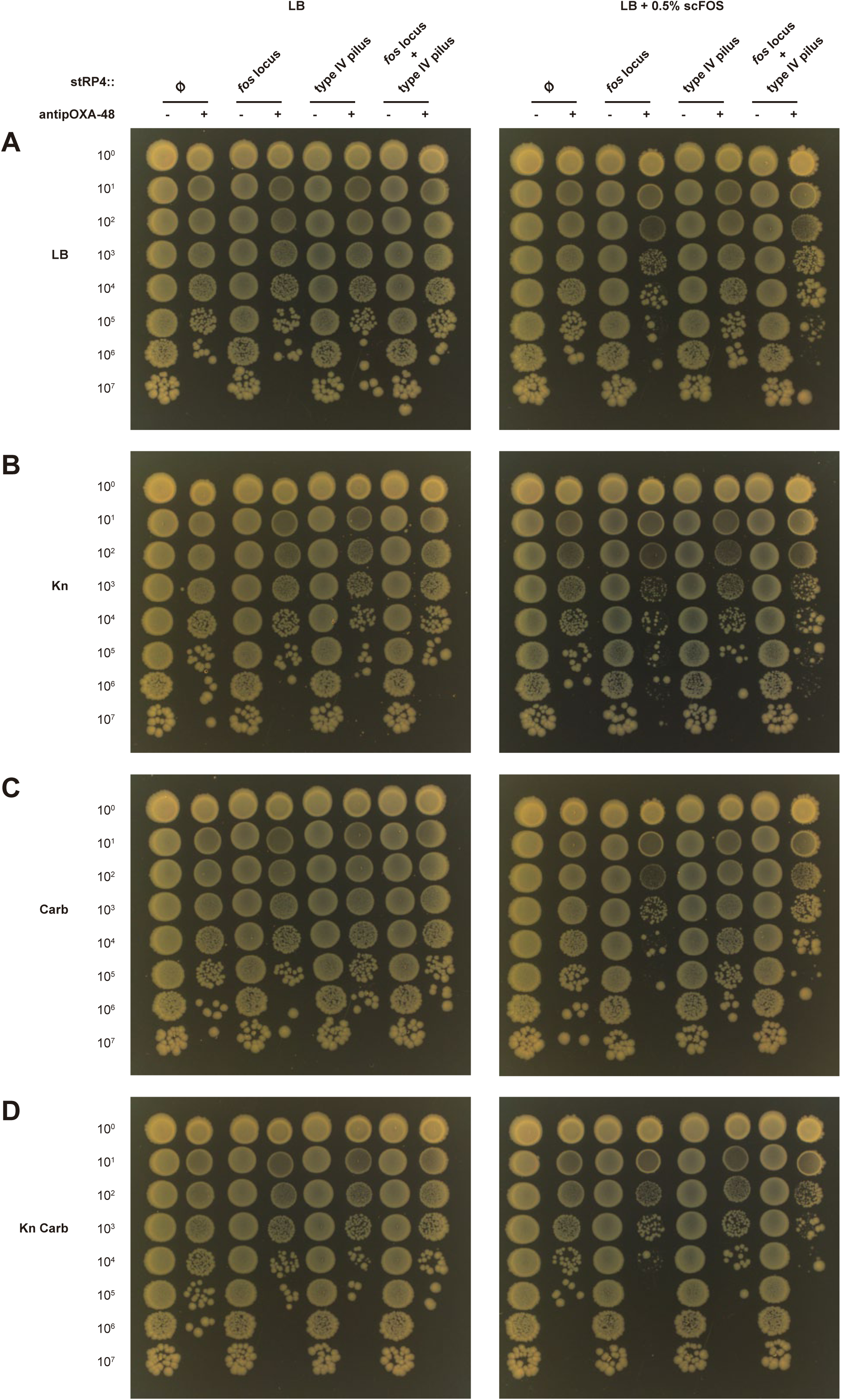
Population-level impact of conjugative delivery of the anti-pOXA-48 toxin–intein module. Population dynamics were assessed after 24 h solid-surface conjugation assays between isogenic *E. coli* populations carrying pOXA-48 and different RP4 derivatives, either lacking or encoding the anti-pOXA-48 toxin–intein module. **Left panels** show the results obtained from conjugations performed in the absence of scFOS, whereas **right panels** correspond to assays conducted in the presence of 0.5% scFOS. Panel **A** depicts total bacterial populations on non-selective medium. Panel **B** shows cells carrying RP4 derivatives (kanamycin selection). Panels **C** shows pOXA-48 carriers (carbenicillin selection). Panel **D** represents cells co-carrying both plasmids (kanamycin and carbapenem selection). The data shown in this figure are representative of at least three independent biological replicates.

**Figure S12.**
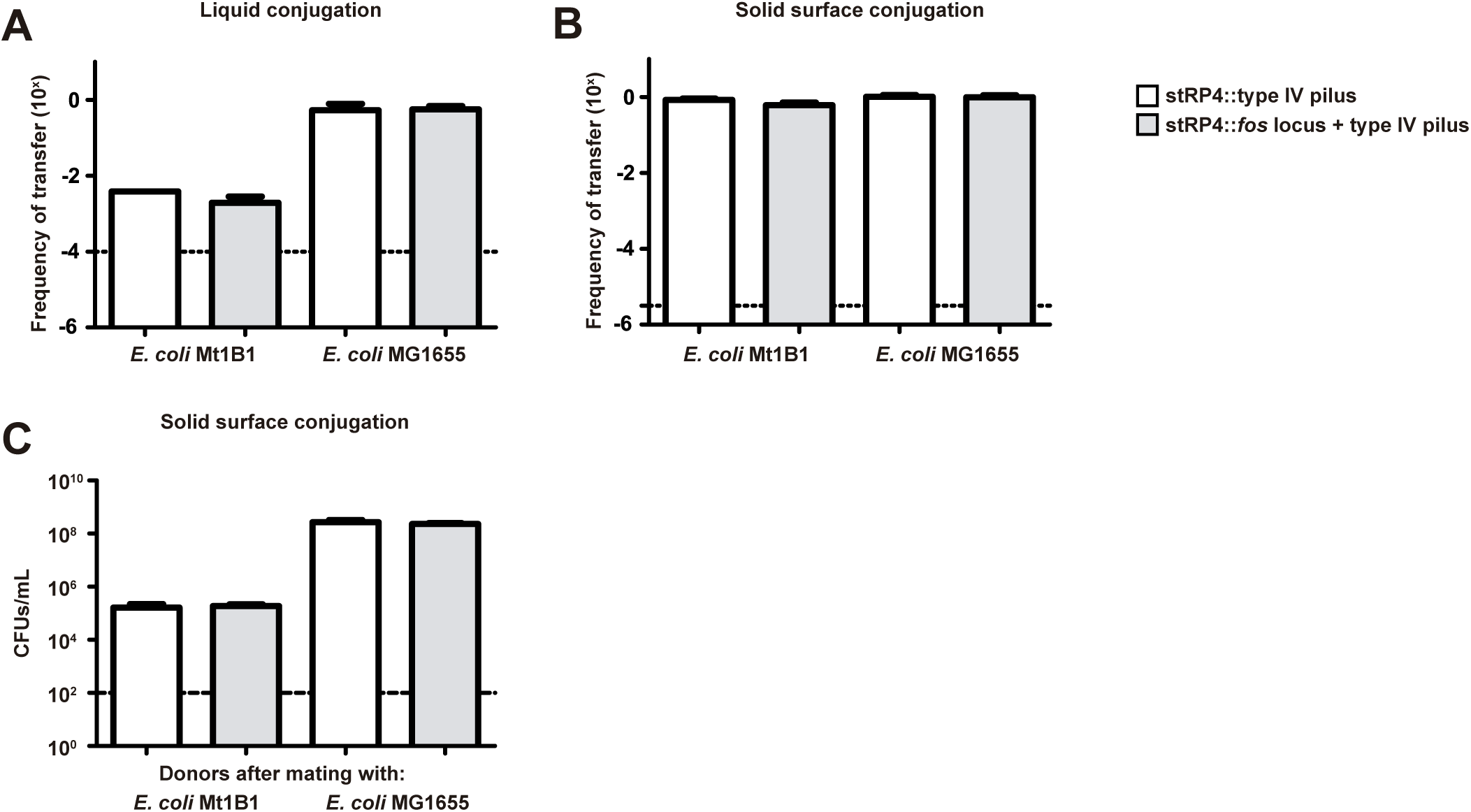
Comparison of transfer efficiencies of stRP4::type IV pilus (white) and stRP4::*fos* locus + type IV pilus (grey) in liquid (**A**) and solid (**B**) conjugation assays. An auxotrophic *E. coli* MG1655 strain was used as the donor, and *E. coli* MG1655 and *E. coli* Mt1B1 were used as recipients. In panel **C**, donor loads quantified after the solid-surface conjugation assays are shown. Data represent mean values from three independent biological replicates. Error bars indicate standard deviation. The dashed line marks the detection limit.

## Supplementary information

### 1. Validation of the *virB* promoter

Given its central regulatory role, VirF represents a promising antivirulence target. For this reason, we sought to exploit this regulatory circuit as the foundation of our anti-*Shigella*/EIEC strategy. As a first step, we evaluated the activity of the *virB* promoter, aiming to assess its responsiveness to VirF and determine its suitability for driving expression of a toxin in a VirF-dependent system. To do this, we fused the *virB* promoter region^1^ to *gfp* (**Figure S2B**) and measured its expression levels at 30°C and 37°C in several *Shigella* species (*S. flexneri*, *S. boydii* and *S. dysenteriae*). *E. coli* MG1655, which lacks pINV, was used as a negative control. This experiment was based on the known thermoregulation of *virF* expression: at temperatures below 32°C, the nucleoid-associated protein H-NS represses *virF* transcription; however, upon entering the human host and experiencing a temperature shift to 37°C, this repression is alleviated, enabling *virF* expression and subsequent activation of the *virB* promoter.

As shown in **Figure S3A**, all *Shigella* species tested exhibited a marked increase in GFP expression at 37°C compared to 30°C, consistent with the well-characterized thermoregulation of *virF*. In contrast, *E. coli* MG1655, which lacks the pINV plasmid, showed no such behaviour.

### 2. Validation of the *hilA* promoter

To determine whether the *hilA* promoter met our criteria for expression strength and specificity, we constructed a reporter system analogous to the one previously developed for *Shigella* replacing the *virB* promoter with that of *hilA*^2^ (**Figure S4B**). As a first step, we transformed *E. coli* MG1655 and *Salmonella enterica* serovar Typhimurium SL1344 with this plasmid and cultured the cells under conditions known to induce *hilD* expression. Specifically, high osmolarity (0.3 M NaCl)^3^ and magnesium levels (10 mM MgSO₄)^4^.

As shown in **Figure S3B**, *E. coli* exhibited negligible GFP production, whereas *Salmonella* displayed a bimodal pattern, with a subpopulation of cells activating the reporter, consistent with previous reports^5,6^. Given the modest signal observed in *Salmonella*, we sought to enhance translation while maintaining specificity. To this end, we modified the native ribosome binding site (RBS) of *hilA* by shortening its distance to the start codon (four nucleotides were removed) and replacing the original sequence (AAGAGAA) with a stronger Shine-Dalgarno motif (AGGAGGA). Upon testing this modified version of the reporter (**Figure S4C**), we saw that the GFP output remained minimal in *E. coli*, whereas in *Salmonella*, the activated subpopulation exhibited a ∼50-fold increase in fluorescence (**Figure S3B**). Therefore, this optimized version of the *hilA* 5’ untranslated region was chosen for the development of our anti-*Salmonella* system.

### 3. Streamlining RP4

With the goal of minimizing plasmid size and eliminating non-essential elements to improve genetic stability and facilitate further engineering, we generated a reduced version of RP4, named stRP4. To achieve this, we removed all sequences not directly involved in the “core” biology of the plasmid, that is, elements known not to affect replication, conjugative transfer, or stability. Specifically, three regions were eliminated (**Figure S8A**):

1. **The IS21 insertion sequence,** which interrupts the aminoglycoside resistance gene *aphA*. In the wildtype RP4 plasmid, despite this disruption, *aphA* remains functional. This is likely due to the IS21 being inserted near the N-terminal end of the gene, which permits the translation of a functional, truncated version of AphA, potentially initiated from an alternative start codon (GTG) and a Shine-Dalgarno-like sequence (GCGCAGCCG) located upstream of it. In addition, the IS must facilitate continued transcription of *aphA* through either an internal promoter within the IS21 element itself or, if no transcriptional terminators are present within the IS, via transcriptional readthrough from upstream. In the reduced version of the plasmid, the uninterrupted *aphA* gene was restored.
2. **The Tn1 transposon,** which harbors the *bla* gene conferring resistance to β-lactam antibiotics, as well as the genes encoding the Tn1 transposase and resolvase. In the wild-type plasmid, Tn1 is inserted within *klcB*. In the reduced version, the entire transposon was removed and the original, uninterrupted *klcB* was restored.
3. **The tetracycline resistance cassette**, from which only the coding sequences, *tetA* and *tetR*, were removed, while the transcriptional terminators were preserved. This region was selected as insertion site for our genetic cargoes, as its relative isolation within the plasmid architecture is expected to minimize unintended transcriptional interference with surrounding elements.

Following assembly of the modified plasmid in *Saccharomyces cerevisiae* and its subsequent transfer into *E. coli*, the plasmid was fully sequenced to verify the absence of unwanted mutations. We then assessed its conjugation frequency and stability under non-selective conditions. These parameters were compared to those of the wildtype plasmid to ensure that the introduced deletions did not compromise essential plasmid functions. As it can be seen in **Figure S8B**, both the wild-type and reduced plasmids remained stably maintained in 100% of the population over a period of 14 days. In terms of conjugation efficiency, the streamlined plasmid displayed comparable transfer rates to those of its wildtype counterpart, and even higher rates in *Salmonella enterica*, as illustrated in the section **C** of the same figure.

### 4. Assessing the functionality of the *fos* locus

After cloning the *fos* locus into the reduced version of the RP4 plasmid, we first confirmed its functionality by assessing the growth of *E. coli* carrying either the stRP4 or its *fos* derivative in rich medium, in presence and absence of exogenously added scFOS. As shown in **Figure S9**, in the absence of supplemented scFOS, the presence of the *fos* locus resulted in a slight increase in overall culture growth. Upon exogenous addition of scFOS, the strain harboring the *fos*-plasmid also reached higher optical densities than those carrying the empty vector. Moreover, in this case, the growth curve of the *fos*-carrying strain displayed a pattern consistent with catabolite repression, suggesting sequential utilization of available carbon sources.

### 5. *E. coli* Mt1B1 conjugation dynamics: limited transfer under liquid conditions and possible involvement of contact-dependent antagonistic systems

To better understand the dynamics observed in the OMM^12^ community experiments (**Figure 5**), we investigated the conjugative behaviour of *E. coli* Mt1B1 in pairwise assays under both liquid and solid mating conditions.

Liquid conjugation assays were performed using auxotrophic MG1655 derivatives carrying stRP4 variants as donors and *E. coli* Mt1B1 as the recipient. Under liquid conditions, transfer frequencies into Mt1B1 were markedly reduced (approximately two orders of magnitude) compared to those observed when MG1655 was used as recipient (**Figure S12A**).

To determine whether this limitation was environment-dependent, conjugation assays were repeated on solid surfaces. Under these conditions, plasmid transfer into Mt1B1 was comparable to that of MG1655 (**Figure S12B**), indicating that physical proximity on surfaces can partially overcome the constraints observed in liquid media. However, when donor loads were quantified at the end of these solid-surface matings (**Figure S12C**), we consistently observed an approximately 3-log reduction in MG1655 donor counts when mated with Mt1B1. This reduction was not detected in control matings in which MG1655 was paired with isogenic MG1655 recipients under the same conditions (**Figure S12C**), suggesting that the effect is specific to interactions with Mt1B1.

These results indicate that Mt1B1 behaves as a poor recipient for conjugation in liquid environments. One possible explanation is impaired mating-pair stabilization. Conjugation mediated by RP4 derivatives equipped with a type IV pilus depends on productive adhesin–recipient interactions, which are influenced by the C-terminal domain of PilV encoded within the shufflon region. The absence of a compatible surface determinant in Mt1B1 could therefore limit stable donor–recipient contacts in liquid culture, restricting plasmid acquisition under these conditions.

On the other hand, the pronounced loss of donor cells during surface-associated mating is consistent with the action of a contact-dependent antagonistic mechanism^7^. One plausible explanation is the activity of a type VI secretion system (T6SS), which mediates interbacterial killing upon close cell–cell contact. Although the involvement of a T6SS was not directly tested in this study, the magnitude of donor reduction and its dependence on surface-associated contact are compatible with such a mechanism.

Overall, under liquid conditions, inefficient plasmid acquisition limits dissemination into Mt1B1, confining plasmid carriage to MG1655-derived populations. On solid surfaces, although transfer can occur, donor fitness is severely compromised, likely due to contact-dependent antagonism. These features provide a mechanistic explanation for the decline of plasmid-bearing populations observed in complex community experiments.

## Notes

### Competing Interest Statement

The authors have declared no competing interest.

